# Evolutionary transcriptomics implicates new genes and pathways in human pregnancy and adverse pregnancy outcomes

**DOI:** 10.1101/2021.04.19.440514

**Authors:** Katie Mika, Mirna Marinić, Manvendra Singh, Vincent J. Lynch

## Abstract

Evolutionary changes in the anatomy and physiology of the female reproductive system underlie the origins and diversification of pregnancy in Eutherian (“Placental”) mammals. This developmental and evolutionary history constrains normal physiological functions and biases the ways in which dysfunction contributes to reproductive trait diseases and adverse pregnancy outcomes. Here, we show that gene expression changes in the human endometrium during pregnancy are associated with the evolution of human-specific traits and pathologies of pregnancy. We found that hundreds of genes gained or lost endometrial expression in the human lineage. Among these are genes that may contribute to human-specific maternal-fetal communication (*HTR2B*) and maternal-fetal immunotolerance (*PDCD1LG2*) systems, as well as vascular remodeling and deep placental invasion (*CORIN*). These data suggest that explicit evolutionary studies of anatomical systems complement traditional methods for characterizing the genetic architecture of disease. We also anticipate our results will advance the emerging synthesis of evolution and medicine (“evolutionary medicine”) and be a starting point for more sophisticated studies of the maternal-fetal interface. Furthermore, the gene expression changes we identified may contribute to the development of diagnostics and interventions for adverse pregnancy outcomes.

## Introduction

Evolutionary changes in the ontogeny of anatomical systems are ultimately responsible for their functional conservation and transformation into new tissue and organ systems (novelties) with new physiological functions that are outside of the range of the ancestral ones (innovations). These same evolutionary and developmental histories limit (constrain) the range of genetic and environmental perturbations those physiological functions can accommodate before leading to dysfunction and disease (i.e., their reaction norms). Evolution of the structures and functions of female reproductive system and extra-embryonic fetal membranes, for example, underlie the evolution of pregnancy (Armstrong et al., 2017; Hou et al., 2009; Kin et al., 2015; Lynch et al., 2015, 2008) and likely adverse pregnancy outcomes such as infertility (Cummins, 1999), recurrent spontaneous abortion (Kosova et al., 2015), preeclampsia (Carter, 2011; Crosley et al., 2013; Elliot, 2017; Rosenberg and Trevathan, 2007), and preterm birth (LaBella et al., 2020; Marinić et al., 2021; Plunkett et al., 2011; Swaggart et al., 2015). Thus, reconstructing the evolutionary and developmental history of the cells, tissues, and organs involved in pregnancy may elucidate the ontogenetic origins and molecular etiologies of adverse pregnancy outcomes.

Extant mammals span major stages in the evolution and diversification of pregnancy, including the origins of maternal provisioning (matrotrophy), placentation, and viviparity (Behringer et al., 2006; Freyer et al., 2003; Freyer and Renfree, 2009; Hughes and Hall, 1998; Renfree, 1995; Renfree and Shaw, 2013). Eutherian mammals have also evolved a complex suite of traits that support prolonged pregnancies such as an interrupted estrous cycle, maternal recognition of pregnancy, maternal-fetal communication, immunotolerance of the antigenically distinct fetus, and implantation of the blastocyst into maternal tissue (Abbot and Rokas, 2017). There is also considerable variation in pregnancy traits within Eutherians. Catarrhine primates, for example, have evolved spontaneous decidualization (differentiation) of endometrial stromal fibroblasts (ESFs) into decidual stromal cells (DSCs) under the combined action of progesterone, cyclic adenosine monophosphate (cAMP), and other unknown maternal signals (Carter and Mess, 2017; Gellersen et al., 2007; Gellersen and Brosens, 2003; Kin et al., 2016, 2015; Mess and Carter, 2006), deeply invasive interstitial hemochorial placentas (Carter et al., 2015; Pijnenborg et al., 2011a, 2011b; Soares et al., 2018), menstruation (Burley, 1979; Emera et al., 2012b; Finn, 1998; Strassmann, 1996), and a unique but unknown parturition signal (Csapo, 1956; Csapo and Pinto-Dantas, 1965). Humans have also evolved longer pregnancy and labor (Bourne, 1970; Keeling and Roberts, 1972), and appear particularly susceptible to pregnancy complications such as preeclampsia (Crosley et al., 2013; Elliot, 2017; Marshall et al., 2018), and preterm birth (Phillips et al., 2015; Rokas et al., 2020; Wildman et al., 2011) than other primates.

Gene expression changes ultimately underlie the evolution of anatomical structures, suggesting that gene expression change at the maternal-fetal interface underlie these primate- and human-specific pregnancy traits. Therefore, we used comparative transcriptomics to reconstruct the evolutionary history of gene expression in the pregnant endometrium and identify genes that gained (“recruited genes”) and lost endometrial expression in the primate and human lineages. We found that genes that evolved to be expressed at the maternal-fetal interface in the human lineage were enriched for immune functions and diseases such as preterm birth and pre-eclampsia. We explored the function of three recruited genes in greater detail, which implicates them in a novel signaling system at the maternal-fetal interface (*HTR2B*), maternal-fetal immunotolerance (*PDCD1LG2*), and remodeling of uterine spiral arteries and deep placental invasion (*CORIN*). These data indicate that explicit evolutionary studies can identify genes and pathways essential for the normal healthy functions cells, tissues, and organs, and that likely underlie the (dys)function of those tissue and organ systems.

## Results

### Endometrial Gene Expression Profiling and Ancestral Transcriptome Reconstruction

To identify gene expression gains and losses in the endometrium that are phylogenetically associated with derived pregnancy traits in humans and Catarrhine primates, we assembled a collection of transcriptomes from the pregnant or gravid endometrium of 20 Eutherian mammals, including human (*Homo sapiens*), baboon (*Papio anubis*), Rhesus monkey (*Macaca mulatta*), and Pig-Tailed macaque (*Macaca nemestrina*), three marsupials, platypus, three birds, and eight lizards, including species that are both oviparous and viviparous (Figure 1A and **Supplementary Table 1**). The complete dataset includes expression information for 21,750 genes and 34 species, which were collected at different gestational times from early-to mid-pregnancy, by multiple labs, and sequencing methods. Thus, differences in transcript abundance between samples may reflect biological differences in mRNA abundances between gestational ages or species, differences in sequencing protocols or other technical factors unrelated to the biology of pregnancy (i.e., batch effects). Therefore, we transformed quantitative gene expression values coded as transcripts per million (TPM) into discrete character states such that genes with TPM ≥ 2.0 were considered expressed (state=1), genes with TPM < 2.0 were considered not expressed (state=0), and missing genes coded as unknown (?) (Box 1). Consistent with significant noise reduction, Multi-Dimensional Scaling (MDS) of species based on gene expression levels (TPMs) was essentially random (Figure 1 – Figure supplement 1A), whereas MDS of the binary encoded dataset grouped species by phylogenetic relatedness (Figure 1 – Figure supplement 1B).

**Figure 1.**
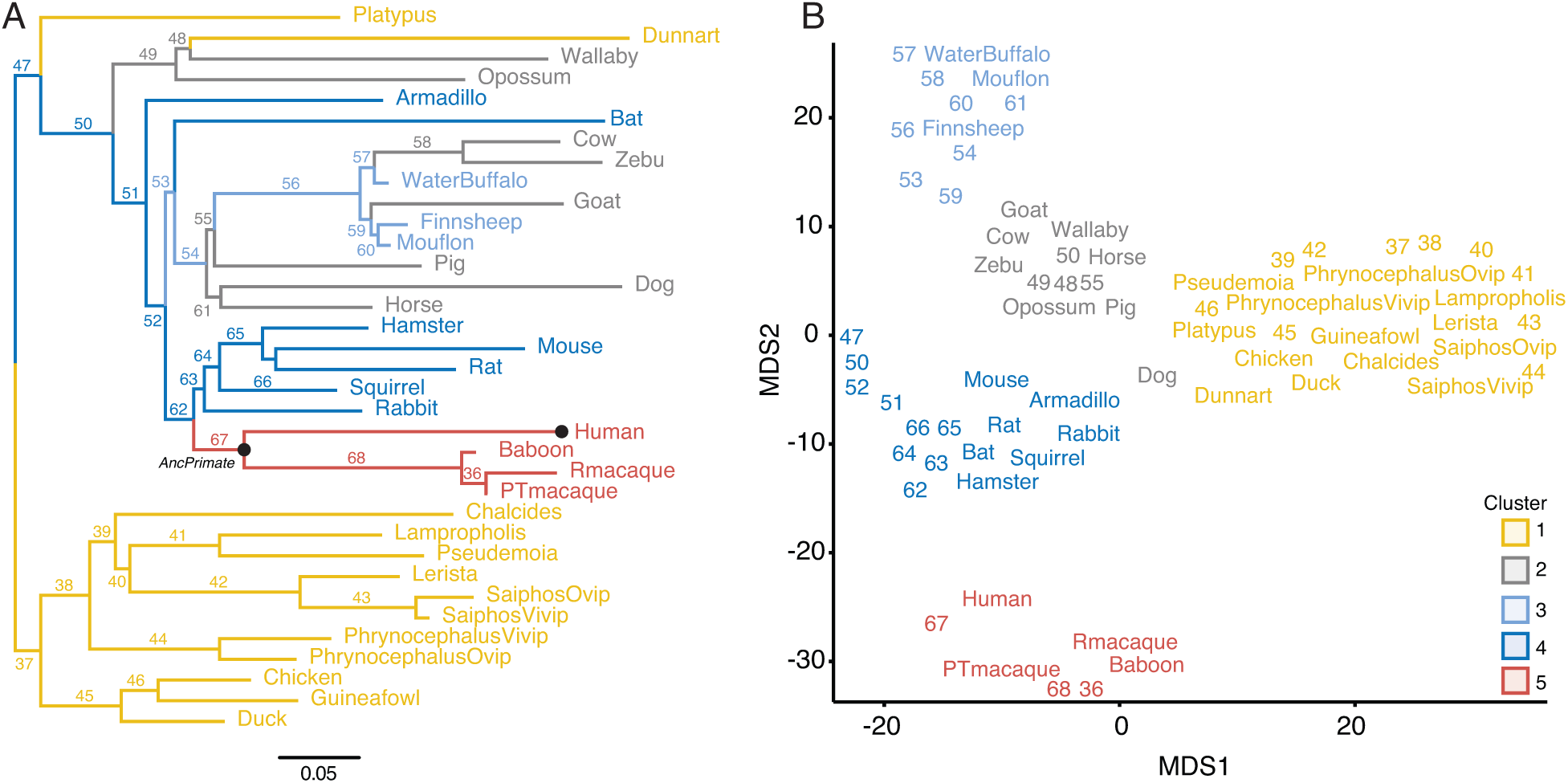
Gene expression profiling at the amniote maternal-fetal interface. **(A)** Amniote phylogeny with branch lengths drawn proportional to the rate of gene expression gains and losses (per gene). The ancestral primate (AncPrimate) and human nodes are indicated with black circles. Internal branches are numbered. **(B)** Multidimensional scaling (MDS) plot of binary encoded endometrial gene expression data for extant and ancestral reconstructed transcriptomes. Transcriptomes are colored by their group membership inferred from K-means clustering with k = 5. Internal branches are numbered.

#### Box 1. Classification of genes into not/expressed categories

A challenge with quantitative gene expression metrics such as RNA-Seq data is defining an expression level that corresponds to functionally active (expressed) genes. Previous studies, however, have shown that an empirically informed operational criterion based on transcript-abundance distributions reasonably approximate gene expression categories (Hebenstreit et al., 2011; Kin et al., 2015; Wagner et al., 2013, 2012). Hebenstreit et al. (2011), for example, showed that genes can be separated into two distinct groups based on their expression levels: the majority of genes follow a normal distribution and are associated with active chromatin marks at their promoters and thus are likely actively expressed, whereas the remaining genes form a shoulder to the left of this main distribution and are unlikely to be actively expressed. Similarly, Wagner et al. (2013, 2012) found that gene expression data could be modeled as a mixture of two distributions corresponding to inactive and actively transcribed genes. Based on this mixture model, they proposed an operational criterion for classifying genes into expressed and non-expressed sets: genes with TPM ≥ 2-4 are likely to be actively transcribed, while genes with TPM < 2 are unlikely to be actively transcribed. Furthermore, Wagner et al. (2013) suggest that the expression cutoff should be chosen depending on the goal of the study. If it is important to reduce false positives (classifying genes as expressed when they are not), then a conservative criterion of TPM ≥ 4 could be used. In contrast, if it is more important to reduce false negative gene expression calls (classifying genes as not expressed when they are), then a liberal criterion such as TPM ≥ 1 could be used. Both Hebenstreit et al. (2011) and Wagner et al. (2013) suggest that genes with TPM ∼ 2 are likely to be actively transcribed.

We found that gene expression data (Log_2_ TPM) from human DSCs generally followed a normal distribution with a distinct shoulder to the left of the main distribution, which could be modeled as a mixture of two Gaussian distributions with means of TPM ∼ 0.11 and TPM ∼ 19 (Box 1 – figure 1A). An empirical cumulative distribution fit (ECDF) to the Gaussian mixture model suggests that genes with TPM = 0.01-1 have less than 50% probability of active expression, whereas genes with TPM ≥ 2 have greater than 75% probability of active expression (Box 1 – figure 1A). Next, we grouped genes into three categories, TPM = 0, TPM = 0.01-1, and TPM ≥ 2, and explored the correlation between these categories and histone marks that are associated with active promoters (H3K4me3) and enhancers (H3K27ac), regions of open chromatin (DNaseI-and FAIRE-Seq), and regions of active transcription (RNA polymerase binding to gene). We found that genes with TPM = 0.01-2 and TPM = 0 were nearly indistinguishable with respect to H3K4me3 marked promoters, H3K27ac marked enhancers, regions of open chromatin assessed by FAIRE-Seq (but not DNaseI-Seq), and most importantly, regions of active transcription as assed by RNA polymerase binding to gene bodies (Box 1 – figure 1B). The promoters of genes with TPM ≥ 2 were also more enriched for binding sites for the progesterone receptor (PGR) and its co-factor GATA2 than genes with TPM < 1 (Box 1 – figure 1C). These data suggest that genes with TPM < 1 are unlikely to be actively expressed while genes with TPM ≥ 2 have hallmarks of active expression. Therefore, we used the TPM ≥ 2.0 cutoff to define a gene as expressed. We note, however, that other cutoffs could be used that either increase or decrease the probability that genes are actively expressed.

**Box 1 – figure 1.**
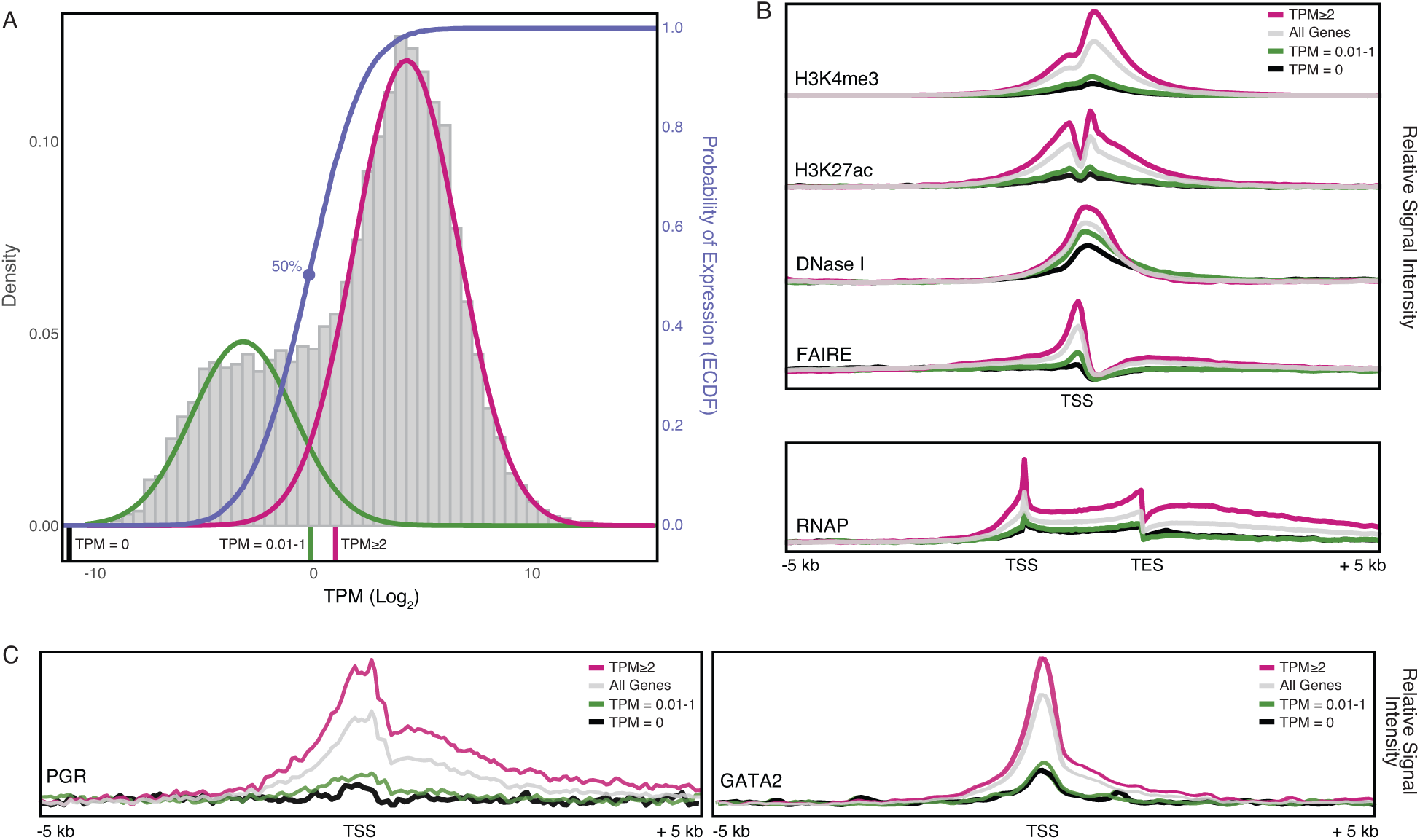
Gene expression and functional genomics data suggest that genes with TPM ≥ 2 are actively expressed. **(A)** Distribution of gene expression levels from human decidual stromal cell (DSC) RNA-Seq data. Gray, kernel density estimates of gene expression levels as transcripts per million (TPM) for human RefSeq genes (genes with TPM = 0 are not shown). Expectation-maximization-based Gaussian mixture curve fits of expression data are shown in green and magenta. Empirical cumulative distribution fit (ECDF) to the Gaussian mixture model is shown in blue. Regions of the kernel density plot corresponding to TPM = 0, TPM = 0.01-1, and TPM ≥ 2 are indicated below the plot as black, green, and pink bars, respectively. The point of the ECDF corresponding to a 50% probability of expression is indicated with a blue circle. **(B)** Correlation of gene expression categories (TPM = 0, TPM = 0.01-1, and TPM ≥ 2) with histone marks that characterize active promoters (H3K4me3), enhancers (H3K27ac), regions of open chromatin (DNaseI and FAIRE), and active transcription (RNAP binding to gene bodies). TSS, transcription start site. TES, transcription end site. **(C)** Correlation of gene expression categories (TPM = 0, TPM = 0.01-1, and TPM ≥ 2) with progesterone receptor (PGR) and the PGR co-factor GATA2 binding sites. TSS, transcription start site.

#### Box 2. Genomic features of human recruited genes

The expression of human recruited genes is enriched in 25 tissues at FDR < 0.05 (Box 2 - figure 1A **inset**), suggesting these genes were predominately recruited into endometrial expression from those tissues. The expression of human recruited genes in human gestation week 9-22 decidua followed a normal distribution that could be modeled as a mixture of two Gaussian distributions with means of TPM ∼2.8 and ∼10.5 (Box 2 – figure 1B), grouping genes into low and high expression sets around TPM ∼4.2 (Box 2 – figure 1B). An empirical cumulative distribution fit (ECDF) to the gene expression data also suggests a cutoff at TPM ∼4.2, which defines an expression level at which 50% of genes are binned into either the high or low expression sets. The promoters of human recruited genes with TPM < 4.2 and ≥ 4.2 were indistinguishable with respect to H3K4me3 and H3K27ac signal at promoters and enhancers, DNaseI hypersensitive sites, PGR, and GATA2 binding, and RNA polymerase binding to gene bodies; both expression sets were generally enriched in these signals compared to genes with TPM = 0 or random genomic locations (Box 2 – figure 1C). In contrast, the promoters of human recruited genes with TPM ≥ 4.2 are in regions of chromatin with greater nucleosome depletion than recruited genes with TPM < 4.2 as assessed by FAIRE-Seq. This observation is consistent with previous studies which found the promoters of highly transcribed genes are preferentially isolated by FAIRE-Seq (Giresi et al., 2007; Nagy et al., 2003).

A particularly noteworthy human recruited gene is *PRL*, which we previously showed evolved endometrial expression in primates (Emera et al., 2012a) and is the most highly expressed human recruited gene in our dataset (Box 2 – figure 1B). Remarkably, *PRL* gene has independently coopted transposable elements (TE) into decidual promoters in multiple Eutherian lineages (Emera et al., 2012a; Gerlo et al., 2006; Lynch et al., 2015), suggesting that TE cooption into decidual promoters may be a widespread phenomenon. Consistent with this hypothesis, human recruited genes with TEs in their promoters and 5’-UTRs had greater H3K4me3 signal and nucleosome depletion as assessed by FAIRE-and DNaseI-Seq (Box 2 – figure 1D). The majority of TEs within the promoters and 5’-UTRs of (40%) were primate-specific, suggesting they may have played a role in recruiting these genes into endometrial expression (Box 2 – figure 1E).

**Box 2 – figure 1.**
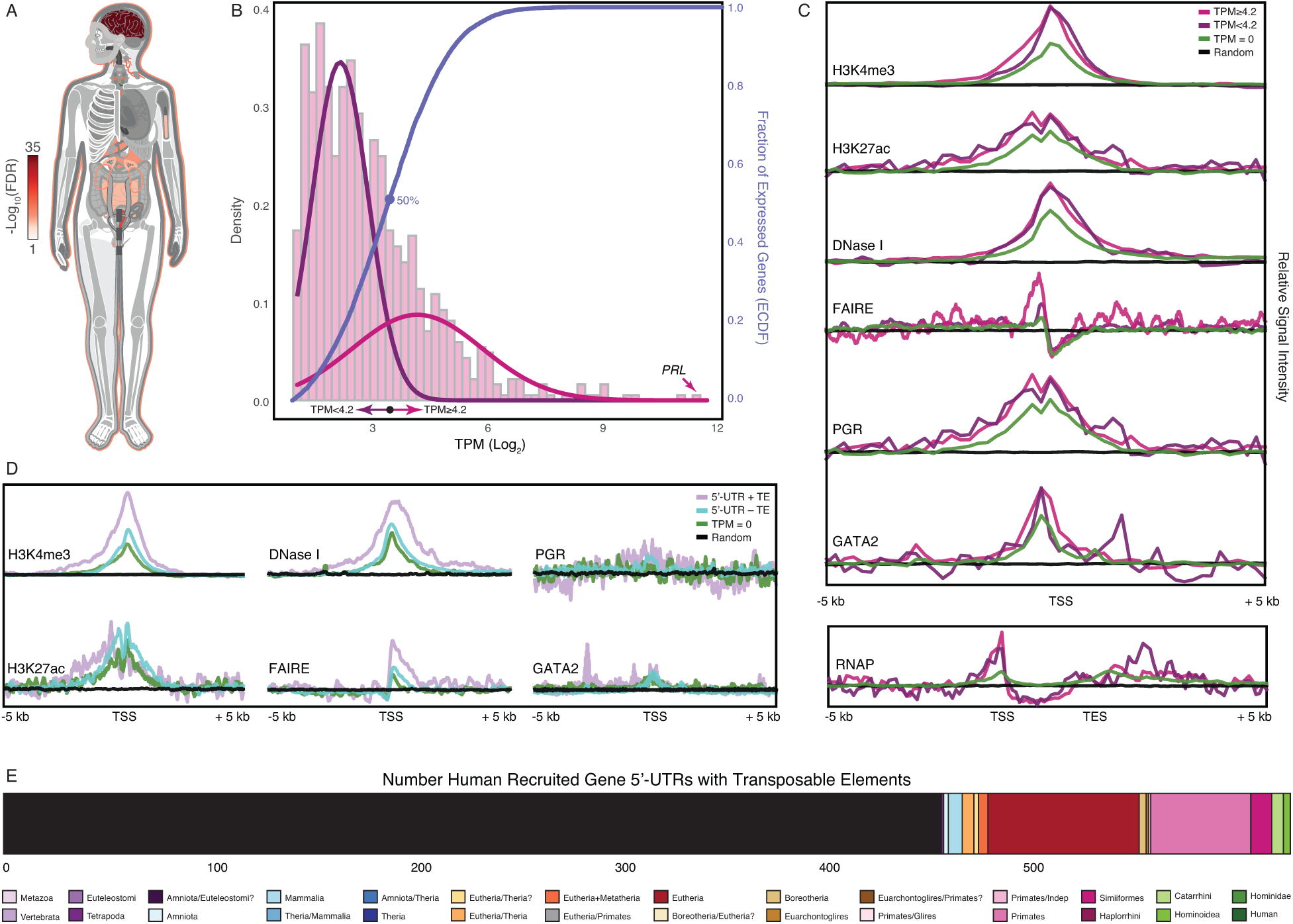
Genomic features of human recruited genes. **(A)** Anatogram heatmap showing organs in which the expression of human recruited genes is enriched (the top 15 organs FDR < 0.05). **(B)** Distribution of human recruited gene expression levels from human gestation week 9-22 decidua RNA-Seq data. Light pink, kernel density estimates of gene expression levels as transcripts per million (TPM) for human RefSeq genes (genes with TPM < 2 are classified as “not expressed” and are not shown). Expectation-maximization-based Gaussian mixture curve fits of expression data are shown in purple (low expressed genes) and magenta (high expressed genes), the TPM 4.2 cutoff for defining low and high expressed genes is shown as a black circle. Empirical cumulative distribution fit (ECDF) to the gene expression data is shown in blue, the point of the ECDF at which 50% of genes are binned into either the high or low expression sets is indicated with a blue circle. **(C)** Correlation of gene expression categories (random genomic locations, TPM = 0, TPM < 4.2, and ≥ 4.2) with histone marks that characterize active promoters (H3K4me3), enhancers (H3K27ac), regions of open chromatin (DNaseI and FAIRE), PGR and GATA2 binding sites, and RNAP binding to gene bodies (active transcription). TSS, transcription start site. TES, transcription end site. **(D)** Correlation of gene expression categories (random genomic locations, TPM = 0, human recruited genes with (+) and without (–) TEs in their promoters) with H3K4me3, H3K27ac, DNaseI, FAIRE, PGR, GATA2, RNAP signal intensities. TSS, transcription start site. TES, transcription end site. **(E)** Number of transposable element families within the promoters and 5’-UTRs of human recruited genes. Transposable elements are colored by their lineage specificity. **Box 2 – source data 1.** Enrichment result table for Figure 1A.

Next, we used the binary encoded dataset to reconstruct ancestral transcriptomes and trace the evolution of gene expression gains (0 à 1) and losses (1 à 0). Ancestral states were inferred with the empirical Bayesian method implemented in IQ-TREE 2 (Minh et al., 2020; Nguyen et al., 2015) using the species phylogeny (Figure 1A) and the GTR2+FO+R4 model of character (Soubrier et al., 2012). Internal branch lengths of the gene expression tree were generally very short while terminal branches were much longer, indicating pronounced species-specific divergence in endometrial gene expression (Figure 1A). MDS of extant and ancestral transcriptomes (Figure 1B) generally grouped species by phylogenetic relationships, parity mode, and degree of placental invasiveness. For example, grouping platypus, birds, and reptiles (cluster 1), viviparous mammals with non-invasive placentas such as opossum, wallaby, and horse, pig, and cow (clusters 2 and 3), and Eutherians with placentas such as mouse, rabbit, and armadillo (cluster 4). Human, baboon, Rhesus monkey, and Pig-Tailed macaque formed a distinct group from other Eutherians (cluster 5), indicating that Catarrhine primates have an endometrial gene expression profile during pregnancy that is distinct even from other Eutherians (Figure 1B).

### Gain and Loss of Signaling and Immune Regulatory genes in Humans

We identified 923 genes that gained endometrial expression in the human lineage with Bayesian Posterior Probabilities (BPP) ≥ 0.80. These genes are enriched in 54 pathways, 102 biological process Gene Ontology (GO) terms, and 91 disease ontologies at a false discovery rate (FDR) ≤ 0.10 (Figure 2). Among enriched pathways were “GPCRs, Class A Rhodopsin-like”, “Signaling by GPCR”, “Cytokine-cytokine receptor interaction”, “Allograft Rejection”, and “Graft-versus-host disease”. The majority of enriched GO terms were related to signaling processes, such as “cAMP-mediated signaling” and “serotonin receptor signaling pathway”’ or to the immune system, such as “acute inflammatory response” and “regulation of immune system process”. The majority of enriched disease ontologies were related to the immune system, such as “Autoimmune Diseases”, “Immune System Disease”, “Inflammation”, “Asthma”, “Rheumatic Diseases”, “Dermatitis”, “Celiac Diseases” and “Organ Transplantation”, as well as “Pregnancy”, “Pregnancy, First Trimester”, “Infertility”, “Habitual Abortion”, “Chorioamnionitis”, “Pre-Eclampsia”, and “Preterm Birth”, consistent with observations that women with systemic autoimmune diseases have an elevated risk of delivering preterm (Kolstad et al., 2019).

**Figure 2.**
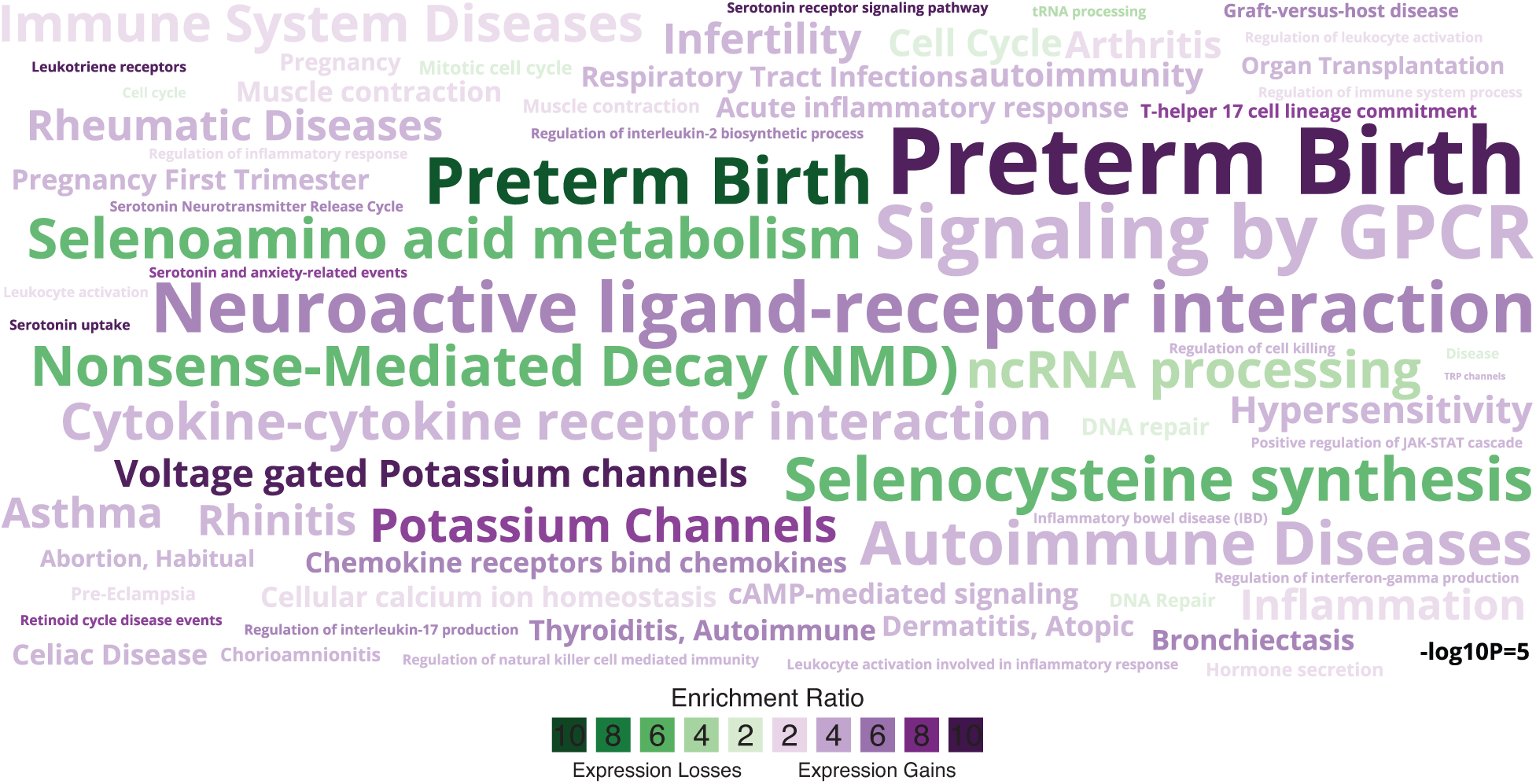
Enriched pathways, gene ontologies, and disease ontologies among genes that gained or lost endometrial expression in the Hominoid (human) lineage. Data shown as a WordCloud, with term size proportional to −log10 hypergeometric P-value (see inset scale) and colored according to enrichment ratio for genes that gained (purple) or lost (green) endometrial expression. **Figure 2 – Source data 1. Custom gmt file used for enrichment tests related to preterm birth.**

771 genes lost endometrial expression in the lineage with BPP ≥0.80. These genes were enriched in 48 pathways, 42 biological process Gene Ontology terms, and 3 disease ontologies at FDR ≤ 0.10 (Figure 2). Enriched pathways included many related to the “immune system”, “pregnancy”, “pregnancy first trimester”, “infertility”, “habitual abortion”, “preeclampsia”, and “preterm birth”. Unlike genes that gained endometrial expression in the human lineage, those that lost endometrial expression were enriched in disease ontologies unrelated to the immune system but did include “Preterm Birth”, as well as “Selenocysteine Synthesis” and “Selenoamino Acid Metabolism”, the latter two which have been previously implicated in preterm birth by GWAS (Zhang et al., 2017). In stark contrast genes that gained (+) or lost (-) endometrial expression during pregnancy in the stem-lineage of Primates (+63/-34) did not include terms related to the immune system or pregnancy. Thus, genes that gained or lost endometrial expression in the human lineage are uniquely related to immune regulatory process, autoimmunity, inflammation, and allograft rejection, signaling processes such as cAMP-mediated and serotonin receptor signaling, and well as adverse pregnancy outcomes.

### Human Recruited Genes Predominantly Remodeled the Transcriptome of Endometrial Stromal Cells

The maternal-fetal interface is composed of numerous maternal and fetal cell-types including endometrial stromal lineage cells (Perivascular, EFSs, and DSCs), uterine natural killer cells (uNK), decidual macrophage (uMP), dendritic cells (DC), T helper cells (Th cells), regulatory T cells (T regs), various innate lymphoid cells (ILCs), and multiple trophoblast cell-types (Suryawanshi et al., 2018; Vento-Tormo et al., 2018; Wang et al., 2020). To infer if genes recruited into endometrial expression in the human lineage are enriched in specific cell-types, we used a previously published single cell RNA-Seq (scRNA-Seq) dataset generated from the first trimester human decidua (Vento-Tormo et al., 2018). to identify cell-types at the maternal-fetal interface (Figure 3A; Figure 3 – figure supplement 1). Next, we determined the observed fraction of human recruited genes expressed in each cell-type compared to the expected fraction, and used a two-way Fisher exact test to identify cell-types that were significantly enriched in human recruited genes. Remarkably, human recruited genes were enriched in five of six endometrial stromal lineage cells, including perivascular endometrial mesenchymal stem cells (pvEMSC) and four populations of DSCs, as well as plasmocytes, endothelial cells, dendritic cells, and extravillus cytotrophoblasts (Figure 3B). Consistent with these observations, the expression of human recruited genes defines distinct cell-types at the maternal-fetal interface (Figure 3 – figure supplement 2).

**Figure 3.**
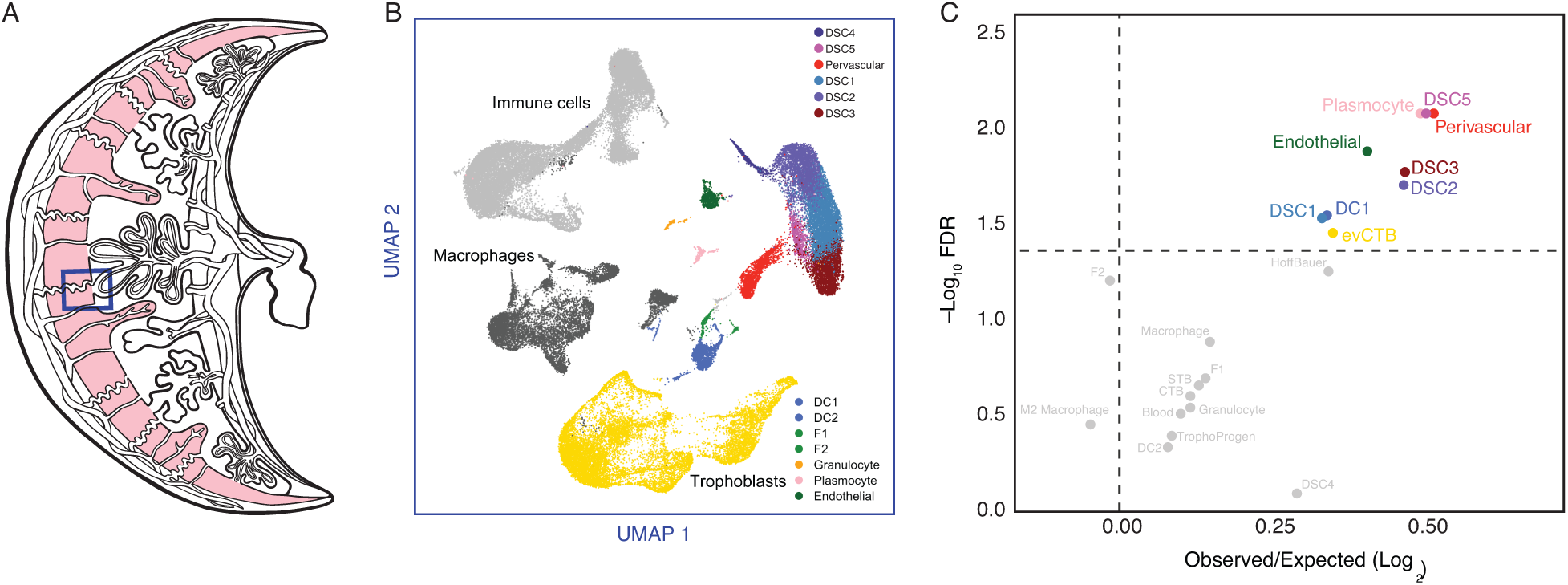
The expression of Hominoid (human) recruited genes enriched in endometrial stromal lineage cells. **(A)** Anatogram of the human maternal-fetal interface. The decidua is shown in light pink, scRNA-Seq data (Vento-Tormo et al., 2018) were generated from the region boxed in blue. **(B)** UMAP clustering of cells from the first trimester maternal-fetal interface. Major cell types and lineages are colored. **(C)** Volcano plot showing cell-types at the maternal-fetal interface in which Hominoid (human) recruited genes are significantly (FDR corrected two-way Fisher’s exact test) enriched (Log_2_ Observed/Expected). Cell-types in which recruited genes are significantly enriched (FDR ≤ 0.05) are labeled and colored as in panel A.

Our observation that human recruited genes have predominantly remodeled the transcriptome of endometrial stromal lineage cells prompted us to explore the development and gene expression evolution of these cell-types in greater detail. Pseudotime single cell trajectory analysis of endometrial stromal lineage cells identified six distinct populations corresponding to a perivascular endometrial mesenchymal stem cell population and five populations of decidual stromal cells (DSC1-5), as well as cells between perivascular endometrial mesenchymal stem cells and DSCs that likely represent non-decidualized endometrial stromal fibroblasts (ESFs) and ESFs that have initiated decidualization (Figure 4A/B). In addition, ESFs that decidualize branch into two distinct lineages, which we term lineage 1 DSCs (DSC1-DSC3) and lineage 2 DSCs (DSC4 and DSC5) (Figure 4A/B). These cell populations differentially express human recruited genes (Figure 4C), which are dynamically expressed during differentiation of perivascular cells into lineage 1 and 2 DSCs (Figure 4D).

**Figure 4.**
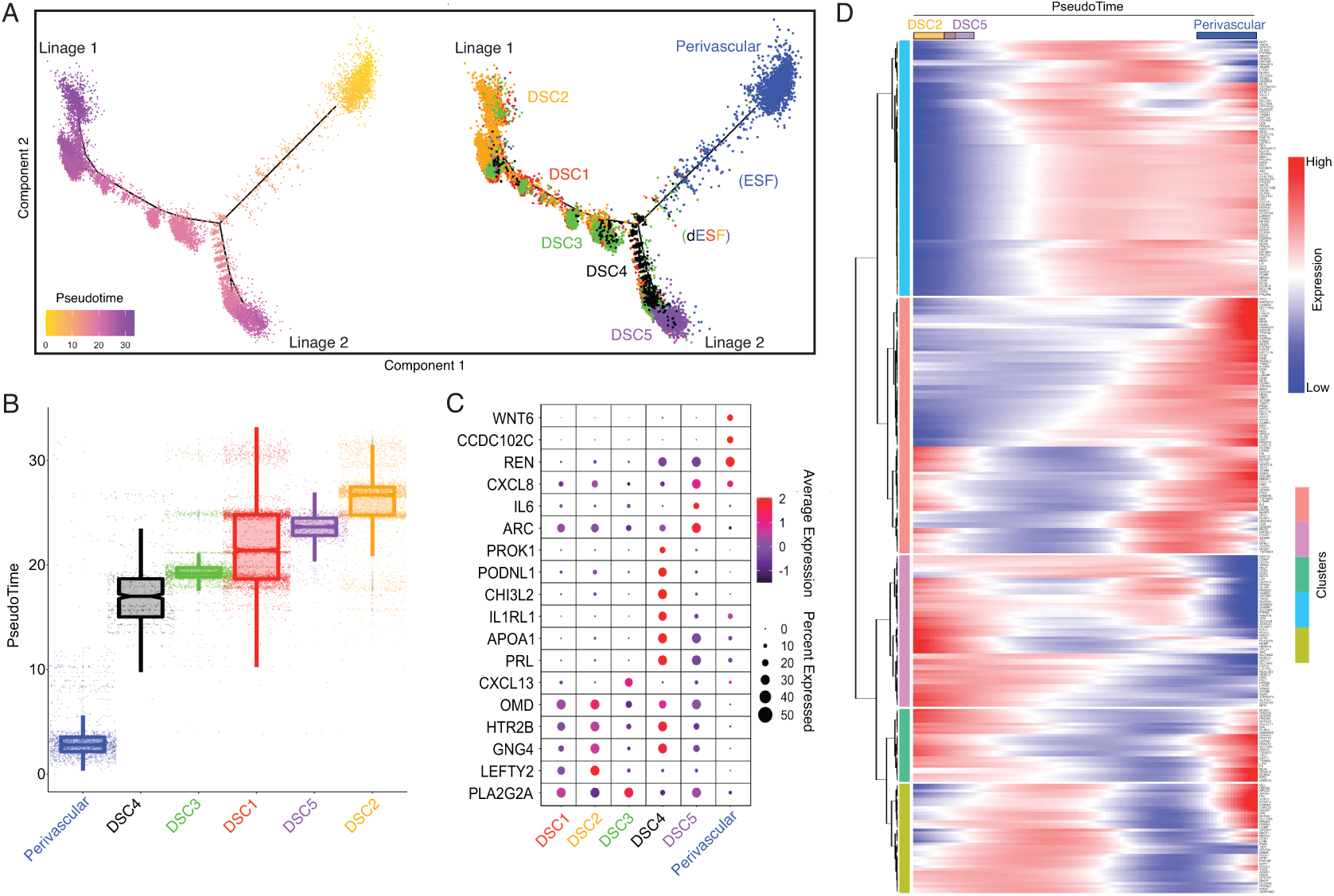
Human-gain genes’ dynamics expression marks the distinct lineages of decidual cells in the spatiotemporal niche. **(A)** Pseudotime trajectory of endometrial stromal lineages, colored from low (gold) to high (purple) pseudotime (left). Endometrial stromal lineage cell-types clustered using the top 2000 differentially expressed genes and projected into a two-dimensional space (right). **(B)** Jittered boxplot illustrating the pseudotime of each cell from A. **(C)** DotPlot illustrating the intensity and abundance of selected human-gain transcript expression between endometrial stromal lineage cell-types. Colors represent an average Log2 expression level scaled to the number of unique molecular identification (UMI) values in single cells. The color scale is from blue to red, corresponding to lower to higher expression, respectively. Dot size is proportional to the percent of cells expressing that gene. Genes were selected based on their differential expression on the pseudotime trajectory shown in the previous figure (Benjamini and Hochberg adjusted P-value < 2.2e-16, Wald test). **(D)** Heatmap showing the kinetics of highly expressed (Log2 scaled average expression > 0.5) human-gain genes changing gradually over the trajectory of endometrial stromal lineage cell-types shown in panel A. Genes (row) are clustered, and cells (column) are ordered according to the pseudotime progression.

### Co-option of Serotonin Signaling in Human Endometrial Cells

Genes that were recruited into endometrial expression in the human lineage are enriched the serotonin signaling pathway (Figure 2), but a role for serotonin signaling in the endometrium has not previously been reported. Among the recruited genes in this pathway is the serotonin receptor *HTR2B*. To explore the history of *HTR2B* expression in the endometrium in greater detail, we plotted extant and ancestral gene expression probabilities on tetrapod phylogeny and found that it independently evolved endometrial expression at least seven times, including in the human lineage (Figure 5A). To investigate which cell-types express *HTR2B,* we used the scRNA-Seq dataset from the first trimester maternal-fetal interface and found that *HTR2B* expression was almost entirely restricted to the DSC cluster (Figure 5B). *HTR2B* was also the only serotonin receptor expressed in either human ESFs or DSCs at TPM ≥ 2 (Figure 5B and Figure 5 – figure supplement 1A) and was highly expressed in uterine tissues (Figure 5 – figure supplement 1B). Additionally, we found that *HTR2B* was only expressed in human and mouse ESFs, but not in ESFs at TPM ≥ 2 from any other tested species (Figure 5C) in a previously generated multispecies ESF RNA-Seq dataset.

**Figure 5.**
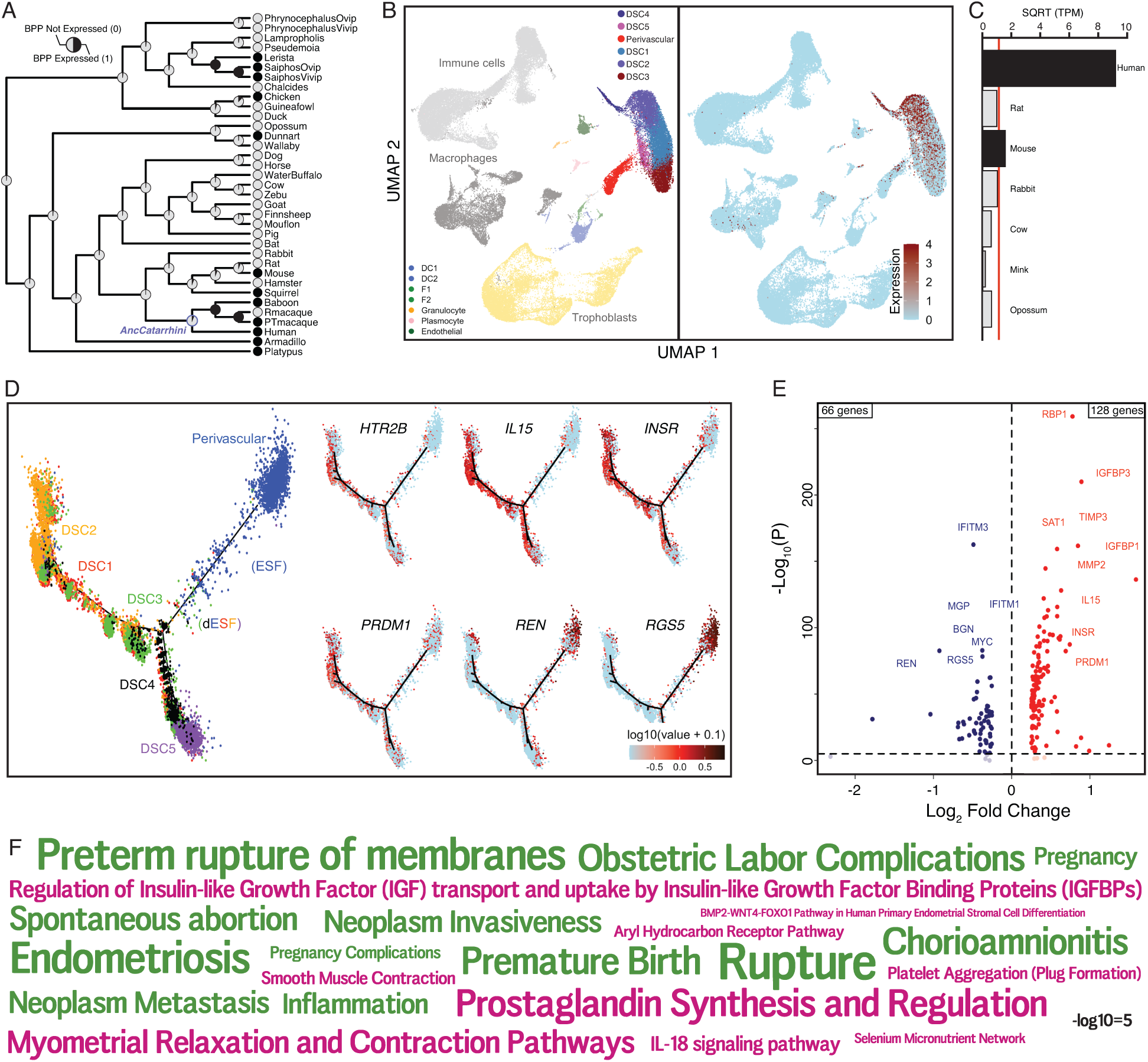
The serotonin receptor *HTR2B* evolved to be expressed in decidual stromal cells at the maternal-fetal interface. **(A)** Ancestral construction of *HTR2B* expression in gravid/pregnant endometrium. Pie charts indicate the Bayesian Posterior Probability (BPP) that *HTR2B* is expressed (state 1) or not expressed (state 0). **(B)** UMAP clustering of cells from the first trimester maternal-fetal interface, decidual stromal cell (DSC) clusters are labeled and highlighted (left). Feature plot based on the UMAP plot showing the single cell expression of *HTR2B* in the endometrial stromal lineage cells. **(C)** Average expression of *HTR2B* in RNA-Seq data from human, rat, mouse, rabbit, cow, mink, and opossum ESFs. Data are shown as square root (SQRT) transformed transcripts per million (TPM), n=2. **(D)** Pseudotime trajectory of endometrial stromal fibroblast lineage cells. Monocle2 visualization of five distinct clusters of DSCs and Perivascular trajectories using the top 2000 differentially expressed genes projected into a two-dimensional space. *HTR2B*, *IL15*, *INSR*, *PRDM1*, *REN*, and *RGS5* expression (log transformed counts) in individual cells are shown in red along the pseudotime trajectory. *IL15*, *INSR*, and *PRDM1* mark DSCs, *REN* and *RGS5* mark Perivascular and decidualizing ESFs (dESFs). **(E)** Volcano plot showing genes that are differentially expressed between *HTR2B^+^ and HTRB^−^* decidual stromal cells. Horizontal dashed line indicates −Log_10_P = 2 (FDR corrected two-way Fisher’s exact test). **(F)** WordCloud showing enriched pathways (pink) and disease ontologies (green) in which genes that are differentially expressed between HTR2B^+^ and HTR2B^-^ cells are enriched.

Pseudotime single cell trajectory analysis of endometrial stromal lineage cells indicates that *HTR2B* is expressed in most lineage 1 DSCs, which co-express other genes such as *IL15*, *INSR*, and *PRDM1* (Figure 5D); *HTR2B* is also expressed by a minority of lineage 2 DSCs, ESFs, and perivascular cells (Figure 5D). 194 genes were differentially expressed between *HTR2B*^+^ and *HTR2B*^−^ DSCs (Figure 5E). These genes were enriched in numerous pathways including “Regulation of Insulin-like Growth Factor (IGF) transport and uptake by Insulin-like Growth Factor Binding Proteins (IGFBPs)”, “Complement and coagulation cascades”, “BMP2-WNT4-FOXO1 Pathway in Human Primary Endometrial Stromal Cell Differentiation”, “IL-18 signaling pathway”, and disease ontologies including “Small-for-dates baby”, “Premature Birth”, “Inflammation”, “Fetal Growth Retardation”, “Pregnancy Complications”, “Hematologic Complications”, and “Spontaneous abortion” (Figure 5F and **Figure 5 – Source data 1**).

To determine if *HTR2B* expression was regulated by progesterone, we used previously published RNA-Seq data from human ESFs and ESFs differentiated into DSCs with cAMP/progesterone (Mazur et al., 2015). *HTR2B* was highly expressed in ESFs and down-regulated during differentiation (decidualization) by cAMP/progesterone into DSCs (Figure 6A and Figure 6 – figure supplement 1). *HTR2B* has hallmarks of an expressed gene in DSCs, including residing in a region open chromatin assessed by previously published FAIRE-Seq data (Figure 6B), an H3K4me3 and H3K27ac marked promoter and polymerase II binding, as well as binding sites for transcription factors that orchestrate decidualization such as the progesterone receptor A isoform (PGR-A), FOXO1, FOSL2, GATA2, and NR2F2 (COUP-TFII) in previously published ChIP-Seq data (see methods) (Figure 6B). The *HTR2B* promoter also makes several long-range interactions to transcription factor bound sites as assessed by H3K27ac HiChIP data generated from a normal hTERT-immortalized endometrial cell line (E6E7hTERT; see methods) (Figure 6B). Consistent with regulation by these transcription factors, knockdown of *PGR*, *FOXO1* and *GATA2* up-regulated *HTR2B* in DSCs (Figure 6C). *HTR2B* is also differentially regulated throughout menstrual cycle (Figure 6D) and pregnancy (Figure 6E; see methods).

**Figure 6.**
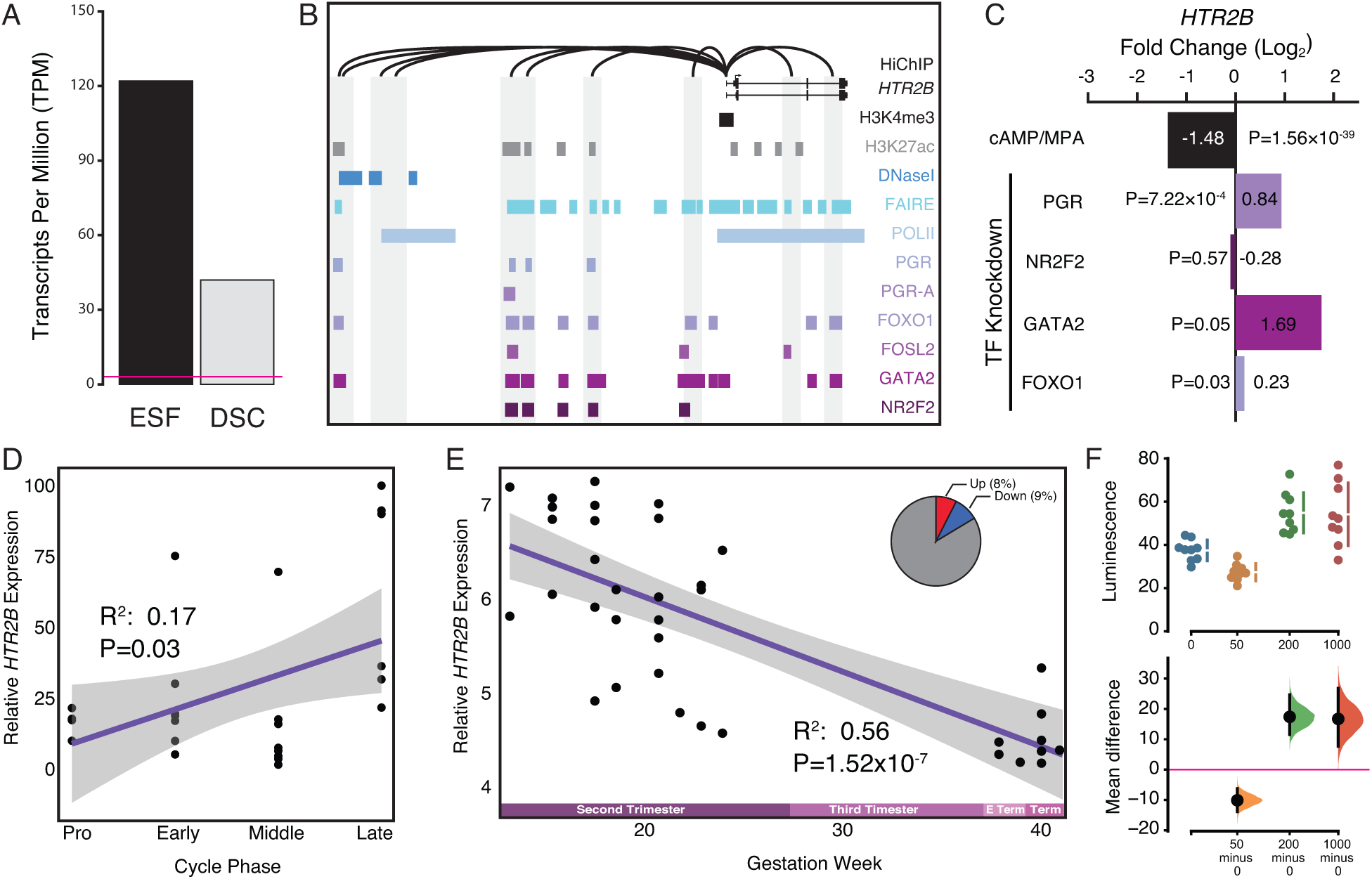
Co-option of serotonin signaling in the endometrium. **(A)** *HTR2B* expression in human ESFs is down-regulated by cAMP/progesterone treatment for 48 hours (decidualization into DSCs). Transcript abundance in RNA-Seq data is shown as transcripts per million (TPM). **(B)** Regulatory elements in human DSCs at the *HTR2B* locus. ChIP-Seq peaks shown for H3K4me3, H3K27ac, polymerase II (POLII), progesterone receptor (PGR) and the PGR-A isoform, FOXO1, FOSL2, GATA2, and NR2F2 (COUP-TFII). Regions of open chromatin are shown from DNaseI-and FAIRE-Seq. Chromatin loops inferred from H3K27ac HiChIP are shown as black arcs connecting the *HTR2B* promoter to other locations in the genome shown in gray. **(C)** *HTR2B* expression is down-regulated by *in vitro* decidualization of ESFs into DSC by cAMP/progesterone treatment, and up-regulated by siRNA-mediated knockdown of PGR, GATA2, and FOXO1, but not NR2F2. n = 3 per transcription factor knockdown. **(D)** Relative expression of HTR2B in the proliferative (n = 6), early (n =4), middle (n = 9), and late (n = 8) secretory phases of the menstrual cycle. Note that outliers are excluded from the figure but not the regression; 95% CI is shown in gray. Gene expression data from Talbi et al., 2006. **(E)** Relative expression of *HTR2B* in the basal plate from mid-gestation to term (14-40 weeks, n = 36); 95% CI is shown in gray. Inset, percent of up-and down-regulated genes between weeks 14-19 and 37-40 of pregnancy (FDR ≤ 0.10). Gene expression data from Winn et al., 2007. **(F)** Cumming estimation plot showing mean difference in luminescence for the serotonin dose response. Upper axis shows relative luminescence of human decidual stromal cells (hDSC) transiently transfected with a luciferase expression vector that drives the transcription of the luciferase reporter gene from a cAMP/PKA response element (pGL4.29[luc2P/CRE/Hygro]) 6 hours after treatment with serotonin (50, 200 and 1000 μM) or vehicle control (VC). Lower axes, mean differences are plotted as bootstrap sampling distributions (n = 5000; the confidence interval is bias-corrected and accelerated). Each mean difference is depicted as a dot. Each 95% confidence interval is indicated by the vertical error bars. P-values indicate the likelihoods of observing the effect sizes, if the null hypothesis of zero difference is true.

To test if human ESFs and DSCs were responsive to serotonin, we transiently transfected each cell-type with reporter vectors that drive luciferase expression in response to activation the AP1 (Ap1_pGL3-Basic[minP]), MAPK/ERK (SRE_pGL3-Basic[minP]), RhoA GTPase (SRF_pGL3-Basic[minP]), and cAMP/PKA (CRE_pGL3-Basic[minP]) signaling pathways, and used a dual luciferase reporter assay to quantify luminescence 6 hours after treatment with either 0, 50, 200, or 1000μM serotonin. Two pathway reporters were responsive to serotonin: 1) The Serum Response Element reporter in DSCs treated with 1000μM serotonin (unpaired mean difference between is 1.35 [95.0%CI 0.624, 2.69], two-sided permutation t-test P = 0.00); and 2) The cAMP/PKA Response Element reporter CRE_pGL3-Basic[minP] in ESFs treated with 1000μM serotonin (unpaired mean difference between is 0.296 [95.0%CI 0.161, 0.43], two-sided permutation t-test P = 0.00) and in DSCs treated with 50μM (unpaired mean difference = −10.1 [95%CI -13.8, -6.28], two-sided permutation t-test P = 0.001), 200μM (unpaired mean difference = 17.4 [95%CI 11.6, 24.6], two-sided permutation t-test P = 0.0004), and 1000μM serotonin (unpaired mean difference is 16.7 [95%CI 7.67, 26.8], two-sided permutation t-test P = 0.006) (Figure 6F and Figure 6 – figure supplement 2).

### Co-option of *PDCD1LG2* (PD-L2) in Human Endometrial Cells

Human recruited genes are enriched numerous immune pathway (Figure 2), among these genes is the PD-1 ligand *PDCD1LG2* (PD-L2) (Figure 7A). We found that *PDCD1LG2* was expressed by several cell-types at the first trimester maternal-fetal interface, including dendritic cells, macrophages, ESFs and DSCs, and multiple trophoblast lineages (Figure 7B), and is highly expressed in uterine tissues (Figure 7 – figure supplement 1). While each of these cell-type populations have individual cells with high level *PDCD1LG2* expression, only 3-5% of DSCs, 3% of dendritic cells, 14% of macrophage, and 66% of cytotrophoblasts express *PDCD1LG2* (Figure 7C). Consistent with recent recruitment in the human lineage, *PDCD1LG2* was highly expressed in human but either moderately or not expressed in ESFs from other species (Figure 7D). The human *PDCD1LG2* locus has the hallmarks of an actively expressed gene, such as a promoter marked by H3K27ac, H3K4me3 and H3K4me1, and binding sites for several transcription factors in previously published ChIP-Seq data from DSCs (Figure 7E). The *PDCD1LG2* promoter also makes several long-range interactions to transcription factor-bound sites, including downstream site that is in the region of open chromatin and bound by PGR/GATA/FOXO1 (Figure 7E). *PDCD1LG2* was highly expressed in ESFs and DSCs (Figure 7F,) but down-regulated by cAMP/progesterone treatment (Figure 7G). Knockdown of *PGR* and *FOXO1* up-regulated and down-regulated *PDCD1LG2* in DSCs, respectively (Figure 7F). *PDCD1LG2* introns also contain several SNPs previously associated with gestational duration and a number of pregnancies in a lifetime, as assessed by GWAS (Aschebrook-Kilfoy et al., 2015; Sakabe et al., 2020; Zhang et al., 2017), albeit with marginal P-values, implicating *PDCD1LG2* in regulating gestation length (Figure 7E).

**Figure 7.**
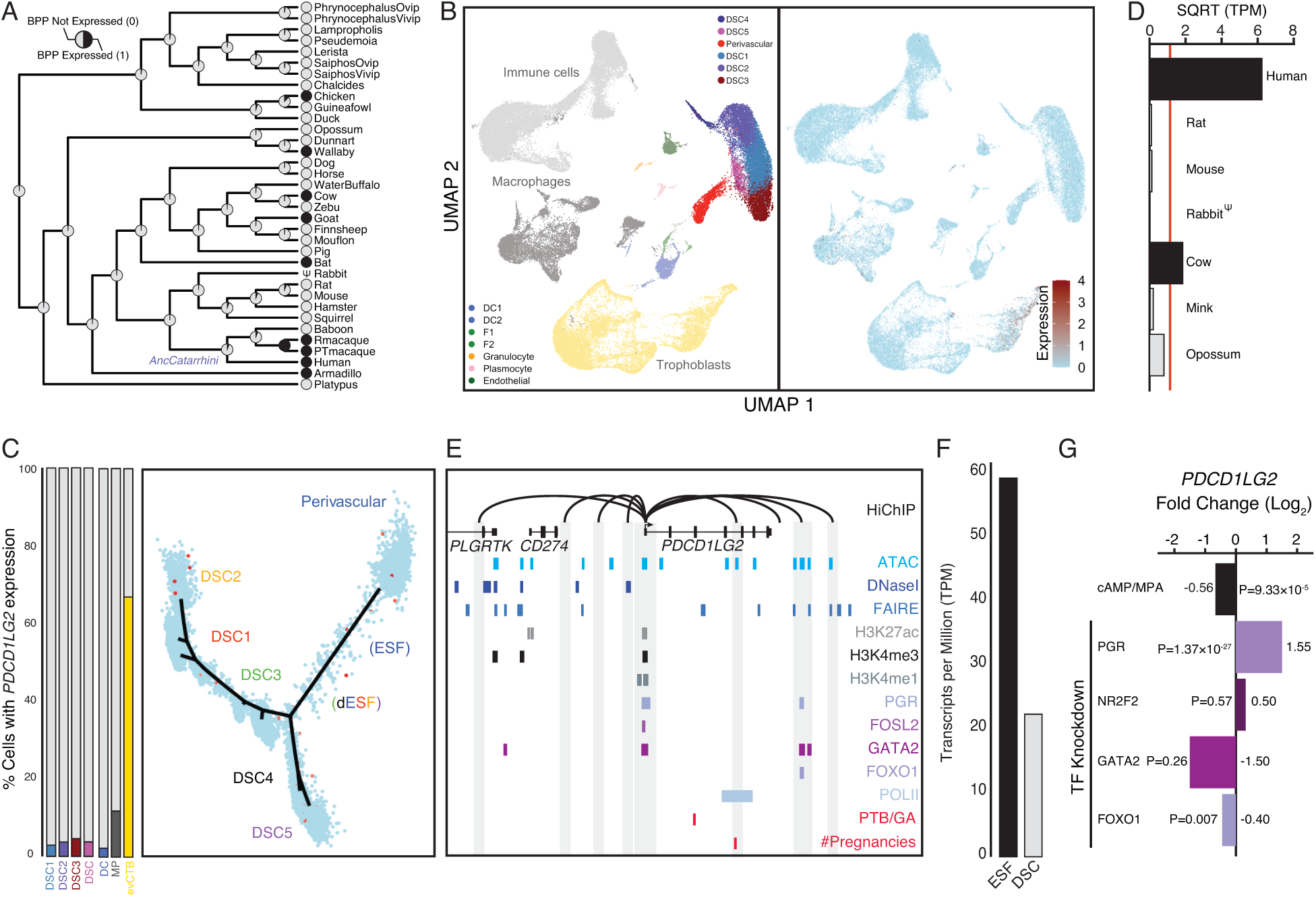
Co-option of *PDCD1LG2* into endometrial cells. **(A)** Ancestral construction of *PDCD1LG2* expression in gravid/pregnant endometrium. Pie charts indicate the Bayesian Posterior Probability (BPP) that *PDCD1LG2* is expressed (state 1) or not expressed (state 0). **(B)** UMAP clustering of cells from the first trimester maternal-fetal interface. *PDCD1LG2* expression (log transformed counts) in individual cells is shown in red. **(C)** Left: Proportion of cell-types at the maternal-fetal interface that express *PDCD1LG2*. Only cell-types that express *PDCD1LG2* are shown as a 100% stacked bar chart: Decidual stromal cell populations 1-3 (DSC1-3), average expression in DSC1-3 (DSC), dendritic cells (DC), macrophage (MP), and extravillus cytotrophoblasts (evCTB). Right: Pseudotime trajectory of endometrial stromal fibroblast lineage cells. Monocle2 visualization of five distinct clusters of DSCs and Perivascular trajectories projected into a two-dimensional space. *PDCD1LG2* expression (log transformed counts) in individual cells is shown in red along the pseudotime trajectory. **(D)** Average expression of *PDCD1LG2* in RNA-Seq data from human, rat, mouse, rabbit, cow, mink, and opossum endometrial stromal fibroblasts (ESFs). Data are shown as square root (SQRT) transformed transcripts per million (TPM), n=2. **(E)** Regulatory elements in human DSCs at the *PDCD1LG2* locus. ChIP-Seq peaks shown for H3K4me1, H3K4me3, H3K27ac, polymerase II (POLII), progesterone receptor (PGR), FOXO1, FOSL2, GATA2, and NR2F2 (COUP-TFII). Regions of open chromatin are shown from DNaseI-, ATAC-and FAIRE-Seq. Chromatin loops inferred from H3K27ac HiChIP are shown as black arcs connecting the *PDCD1LG2* promoter to other locations in the genome shown in gray. The location of SNPs implicated by GWAS in preterm birth is shown in red. **(F)** *PDCD1LG2* expression in human ESFs is down-regulated by cAMP/progesterone treatment for 48 hours (decidualization into DSCs). Transcript abundance in RNA-Seq data is shown as transcripts per million (TPM). **(G)** *PDCD1LG2* expression is down-regulated by *in vitro* decidualization of ESFs into DSC by cAMP/progesterone treatment and by siRNA-mediated knockdown of FOXO1. siRNA-mediated knockdown of PGR up-regulated *PDCD1LG2* expression, while there was no effect after siRNA-mediated knockdown of NR2F2 or GATA2. n = 3 per transcription factor knockdown.

### Co-option of *CORIN* into Human Endometrial Cells

Among the human recruited genes enriched in disease ontologies related to pre-eclampsia (Figure 2) is *CORIN* (Figure 8A), a serine protease which promotes uterine spiral artery remodeling and trophoblast invasion. We found that *CORIN* was exclusively expressed by a subset of endometrial stromal lineage cells (Figure 8B/C), dramatically up-regulated in DSCs by cAMP/progesterone treatment (Figure 8C), and highly expressed in uterine tissues (Figure 8-figure supplement 1). The *CORIN* locus has hallmarks of an actively expressed gene in DSCs, including a promoter in a region of open chromatin assessed by previously published ATAC-and DNase-Seq data and marked by H3K4me3 in previously published ChIP-Seq data (Figure 8E). The *CORIN* promoter also makes long-range interactions to transcription factor-bound sites as assessed by HiChIP, including an up-stream site bound by PGR, FOSL2, GATA2, FOXO1, and NR2F2 in previously published ChIP-Seq data from DSCs (Figure 8E). Consistent with these observations, knockdown of PGR, NR2F2, and GATA2 down-regulated *CORIN* expression in DSCs (Figure 8F).

**Figure 8.**
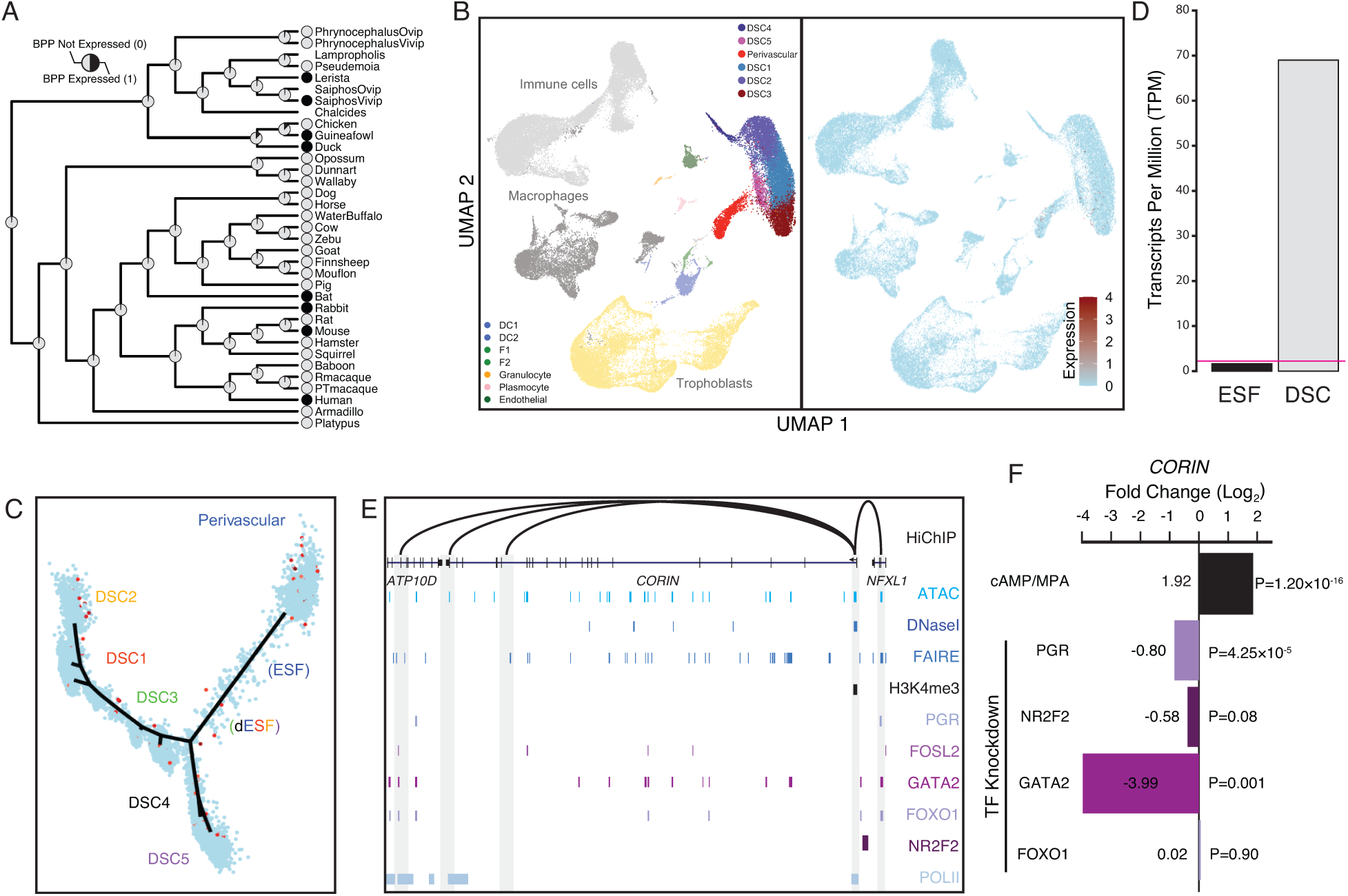
Co-option of *CORIN* into endometrial cells. **(A)** Ancestral construction of *CORIN* expression in gravid/pregnant endometrium. Pie charts indicate the Bayesian Posterior Probability (BPP) that *CORIN* is expressed (state 1) or not expressed (state 0). **(B)** UMAP clustering of cells from the first trimester maternal-fetal interface. *CORIN* expression (log transformed counts) in individual cells is shown in red. **(C)** Pseudotime trajectory of endometrial stromal fibroblast lineage cells. Monocle2 visualization of five distinct clusters of DSCs and Perivascular trajectories projected into a two-dimensional space. *CORIN* expression (log transformed counts) in individual cells is shown in red along the pseudotime trajectory. **(D)** *CORIN* expression in human ESFs is up-regulated by cAMP/progesterone treatment for 48 hours (decidualization into DSCs). Transcript abundance in RNA-Seq data is shown as transcripts per million (TPM). **(E)** Regulatory elements in human DSCs at the *CORIN* locus. ChIP-Seq peaks shown for H3K4me3, polymerase II (POLII), progesterone receptor (PGR), FOXO1, FOSL2, GATA2, and NR2F2 (COUP-TFII). Regions of open chromatin are shown from DNaseI-, ATAC-and FAIRE-Seq. Chromatin loops inferred from H3K27ac HiChIP are shown as black arcs connecting the *CORIN* promoter to other locations in the genome shown in gray. **(F)** *CORIN* expression is up-regulated by *in vitro* decidualization of ESFs into DSC by cAMP/progesterone treatment, and down-regulated by siRNA-mediated knockdown of PGR and GATA2, but not FOXO1 or NR2F2. n = 3 per transcription factor knockdown.

## Discussion

Reconstructing the developmental and evolutionary history of anatomical systems is essential for a causally complete explanation for the origins and progression of disease, which has led to the synthesis of evolution and medicine (“evolutionary medicine”) (Benton et al., 2021). We used comparative transcriptomics to explore how the functions of the maternal side (endometrium) of the maternal-fetal interface evolved, and found that hundreds of genes gained or lost endometrial expression in the human lineage. These recruited genes are enriched in immune functions, signaling processes and genes associated with adverse pregnancy outcomes such as infertility, recurrent spontaneous abortion, pre-eclampsia, and preterm birth. Among these genes are those that may contribute to a previously unknown maternal-fetal communication system (*HTR2B*), augment maternal-fetal immunotolerance (*PDCD1LG2* also known as *PD-L2*), and promote vascular remodeling and deep placental invasion (*CORIN*).

### Human-Specific Remodeling of the Endometrial Stromal Cell Transcriptome

The maternal-fetal interface is composed of numerous maternal cell-types, all which could have been equally impacted by genes that were recruited into endometrial expression in the human lineage. It is notable, therefore, that the expression of these genes is predominately enriched in endometrial stromal lineage cells, including perivascular mesenchymal stem-cells and multiple populations of decidual stromal cells. These data suggest that remodeling of the transcriptome and functions of the endometrial stromal cell lineage has played a particularly important role in the evolution of human-specific pregnancy traits. It is also interesting to note that decidual stromal cells evolved in the stem-lineage of Eutherian mammals (Carter and Mess, 2017; Gellersen et al., 2007; Gellersen and Brosens, 2003; Kin et al., 2016, 2015; Mess and Carter, 2006), coincident with a wave of gene expression recruitments and losses that also dramatically remodeled their transcriptomes (Kin et al., 2015; Lynch et al., 2015). Thus, the endometrial stromal cell lineage has repeatedly been the target of evolutionary changes related to pregnancy, highlighting the importance of decidual stromal cells in the origins and divergence of pregnancy traits. These data also suggest that endometrial stromal lineage cells may play a dominant role in the ontogenesis of adverse pregnancy outcomes.

### Co-option of Serotonin Signaling in Human Endometrial Cells

Unexpectedly, human recruited genes are enriched in the serotonin signaling pathway, such as the serotonin receptor *HTR2B*. Though a role for serotonin in the endometrium has not previously been reported, we found that serotonin treatment effected RAS/MAPK(ERK) and cAMP/PKA signaling pathways, which are essential for decidualization, and that *HTR2B* is dynamically expressed during menstrual cycle and pregnancy, reaching a low at term. Previous studies have shown that the human placenta is a source of serotonin throughout gestation (Clark et al., 1980; Kliman et al., 2018; Laurent et al., 2017; Ranzil et al., 2019; Rosenfeld, 2019). Remarkably, a body of early literature suggests serotonin might trigger parturition. For example, levels of both serotonin (5-HT) and the main metabolite of serotonin (5-HIAA) are highest in amniotic fluid near term and during labor (Jones and Pycock, 1978; Koren et al., 1961; Loose and Paterson, 1966; Tu and Wong, 1976) while placental monoamine oxidase activity (which metabolizes serotonin) is lowest at term (Koren et al., 1965). Furthermore, a single dose of the monoamine oxidase inhibitor paraglyline hydrochloride can induced abortion in humans and other animals (Koren et al., 1966) Consistent with a potential role in regulating gestation length and parturition, use of selective serotonin reuptake inhibitors (SSRIs) is associated with preterm birth (Eke et al., 2016; Grzeskowiak et al., 2012; Huybrechts et al., 2014; Ross et al., 2013; Sujan et al., 2017; Yonkers et al., 2012). 5-HIAA also inhibits RAS/MAPK signaling, potentially by competing with serotonin for binding sites on serotonin receptors (Chen et al., 2011; Klein et al., 2018; Schmid et al., 2015). Collectively, these data suggest a mechanistic connection between serotonin/5-HTAA, and the establishment, maintenance, and cessation of pregnancy.

### Co-option of *PDCD1LG2* (PD-L2) into Human Endometrial Cells

Among the genes with immune regulatory roles that evolved endometrial expression in the human lineage is the programmed cell death protein 1 (PD-1) ligand *PDCD1LG2*. PD-1, a member of the immunoglobulin superfamily expressed on T-cells and pro-B cells, regulates a critical immune checkpoint that plays an essential role in down-regulating immune responses and promoting self-tolerance by suppressing T-cell inflammatory activity (Patsoukis et al., 2020). PD-1 has two ligands, *CD274* (PD-L1) and *PDCD1LG2* (PD-L2), which upon binding PD-1 promote apoptosis in antigen-specific T-cells and inhibit apoptosis in anti-inflammatory regulatory T-cells (Patsoukis et al., 2020). Unlike *CD274*, which is constitutively expressed at low levels in numerous cell-types and induced by IFN-gamma, *PDCD1LG2* expression is generally restricted to professional antigen presenting cells such as dendritic cells and macrophages and has a 4-fold stronger affinity for PD-1 than does *CD274* (Ghiotto et al., 2010; Latchman et al., 2001; Sharpe et al., 2007; Sharpe and Pauken, 2018). These data suggest that a sub-population of human DSCs have co-opted some of the immune regulatory functions of professional antigen presenting cells (APCs), which may have been significantly augmented in the human lineage. While more mechanistic studies will help define the precise role of decidual cells in the establishment and maintenance of maternal-fetal immunotolerance, a role for decidual *PDCD1LG2* in pregnancy is strongly suggested by its association with variants linked to gestational length and number of lifetime pregnancies (parity).

### Co-option of *CORIN* into Human Endometrial Cells

Placental invasiveness varies dramatically in Eutherians, but the cellular and molecular mechanisms responsible for this variation are ill defined. One of the genes that may play a role in the evolution of deeply invasive hemochorial placentation is the serine protease *CORIN*, which converts pro-atrial natriuretic peptide (pro-ANP) to biologically active ANP (Yan et al., 2000). CORIN-mediated ANP production in the uterus during pregnancy has been shown to promote spiral artery remodeling and trophoblast invasion(Cui et al., 2012). These data implicate co-option of *CORIN* into endometrial expression may have contributed to the evolution of particularly deep trophoblast invasion and extensive spiral artery remodeling in humans and other great apes (Carter et al., 2015; Pijnenborg et al., 2011a, 2011b; Soares et al., 2018). *CORIN* expression is also significantly lower in patients with preeclampsia than in normal pregnancies (Cui et al., 2012), suggesting that co-option of *CORIN* into human endometrium may predispose humans to preeclampsia. Additional evolutionary and molecular studies will be required to establish a mechanistic connection between the co-option of *CORIN* into the endometrium, the evolution of hemochorial placentation, and the origins of preeclampsia in the human lineage.

### Caveats and Limitations

A limitation of this study is our inability to determine with precise phylogenetic resolution the lineages in which some gene expression changes occurred. For example, we lack pregnant endometrial samples from Hominoids other than humans (chimpanzee/bonobo, gorilla, orangutan, and gibbon/siamang), thus we are unable to identify truly human-specific gene expression changes. Similarly, we lack endometrial gene expression data from multiple human populations exposed to differing environmental stresses, and therefore are unable to determine the range of physiologically “normal” gene expression or the reaction norms of individual and collective gene expression levels. Our functional genomic and experimental studies are also restricted to an *in vitro* cell culture system, which makes it difficult to assess the *in vivo* impact of gene expression changes on the biology of pregnancy. These limitations are not unique to our study and impact virtually all investigations of Hominoid development and disease, particularly the ones of human-specific traits. Endometrial organoids and iPSC-derived endometrial stromal fibroblasts, however, are promising systems in which to study the development of these traits and disease susceptibility that circumvents the limitations of studying human biology (Abbas et al., 2020; Boretto et al., 2017; Marinić et al., 2020; Turco et al., 2017).

Our gene expression dataset also represents only a snapshot in time of gestation, rather than a comprehensive time course of endometrial gene expression throughout gestation. Interestingly however, the expression changes we identified from these early time points are enriched in disease ontology terms related to adverse pregnancy outcomes that span the length of gestation including infertility, recurrent spontaneous abortion, pre-eclampsia, and preterm birth. These findings suggest that atypical gene expression patterns and physiological changes at the earliest stages, perhaps even processes occurring in the endometrium before pregnancy (e.g., decidualization of ESFs into DSCs), may predispose to multiple adverse outcomes, including those at the latter stages like preterm birth (birth before 37 weeks). An important focus of future studies should be collecting endometrial samples across species and from multiple stages of pregnancy, particularly close to term, when the mechanisms that maintain gestation cease and those that initiate parturition are likely to be activated.

## Conclusions

We found that hundreds of genes gained or lost endometrial expression in humans, including genes that may contribute to a previously unknown maternal-fetal communication system (*HTR2B*), enhanced mechanisms for maternal-fetal immunotolerance (*PDCD1LG2* also known as *PD-L2*), and deep placental invasion (*CORIN*). These results demonstrate that gene expression changes at the maternal-fetal interface likely underlie human-specific pregnancy traits and adverse pregnancy outcomes. Our work also illustrates the importance of evolutionary studies for investigating human-specific traits and diseases. This “evolutionary forward genomics” approach complements traditional forward and reverse genetics in model organisms, which may not be relevant in humans, as well as commonly used methods for characterizing the genetic architecture of disease, such as quantitative trait mapping and genome wide association studies (GWAS). Specifically, our data demonstrate the importance of evolutionary medicine for a mechanistic understanding of endometrial (dys)function, and suggest that similar studies of other tissue and organ systems will help identify genes underlying normal and pathological anatomy and physiology. We anticipate that our results will further the synthesis of evolution and medicine and may contribute to the development of interventions for adverse pregnancy outcomes such as preterm birth.

## Materials and Methods

### Endometrial Gene Expression Profiling

Anatomical terms referring to the glandular portion of the female reproductive tract (FRT) specialized for maternal-fetal interactions or shell formation are not standardized. Therefore, we searched the NCBI BioSample, Sequence Read Archive (SRA), and Gene Expression Omnibus (GEO) databases using the search terms “uterus”, “endometrium”, “decidua”, “oviduct”, and “shell gland” followed by manual curation to identify those datasets that included the region of the FRT specialized for maternal-fetal interaction or shell formation. Datasets that did not indicate whether samples were from pregnant or gravid females were excluded, as were those composed of multiple tissue types. For all RNA-Seq analyses we used Kallisto (Bray et al., 2016) version 0.42.4 to pseudo-align the raw RNA-Seq reads to reference transcriptomes (see **Figure 1 – Source data 1** for accession numbers and reference genome assemblies) and to generate transcript abundance estimates. We used default parameters bias correction, and 100 bootstrap replicates. Kallisto outputs consist of transcript abundance estimates in Transcripts Per Million (TPM), which were used to determine gene expression. To ensure that human decidua samples were free from trophoblast contamination, we compared the expression of placental enriched genes in RNA-Seq data from human placenta, a human endometrial stromal fibroblast (ESF) cell line, a human decidual stromal (DSC) cell line, and human first trimester decidua. These results suggest that there likely no trophoblast contamination of human first trimester decidua samples (Table 1), thus inferences of gene expression gains in the human lineage are unlikely to be the result of trophoblast contamination.

**Table 1.**
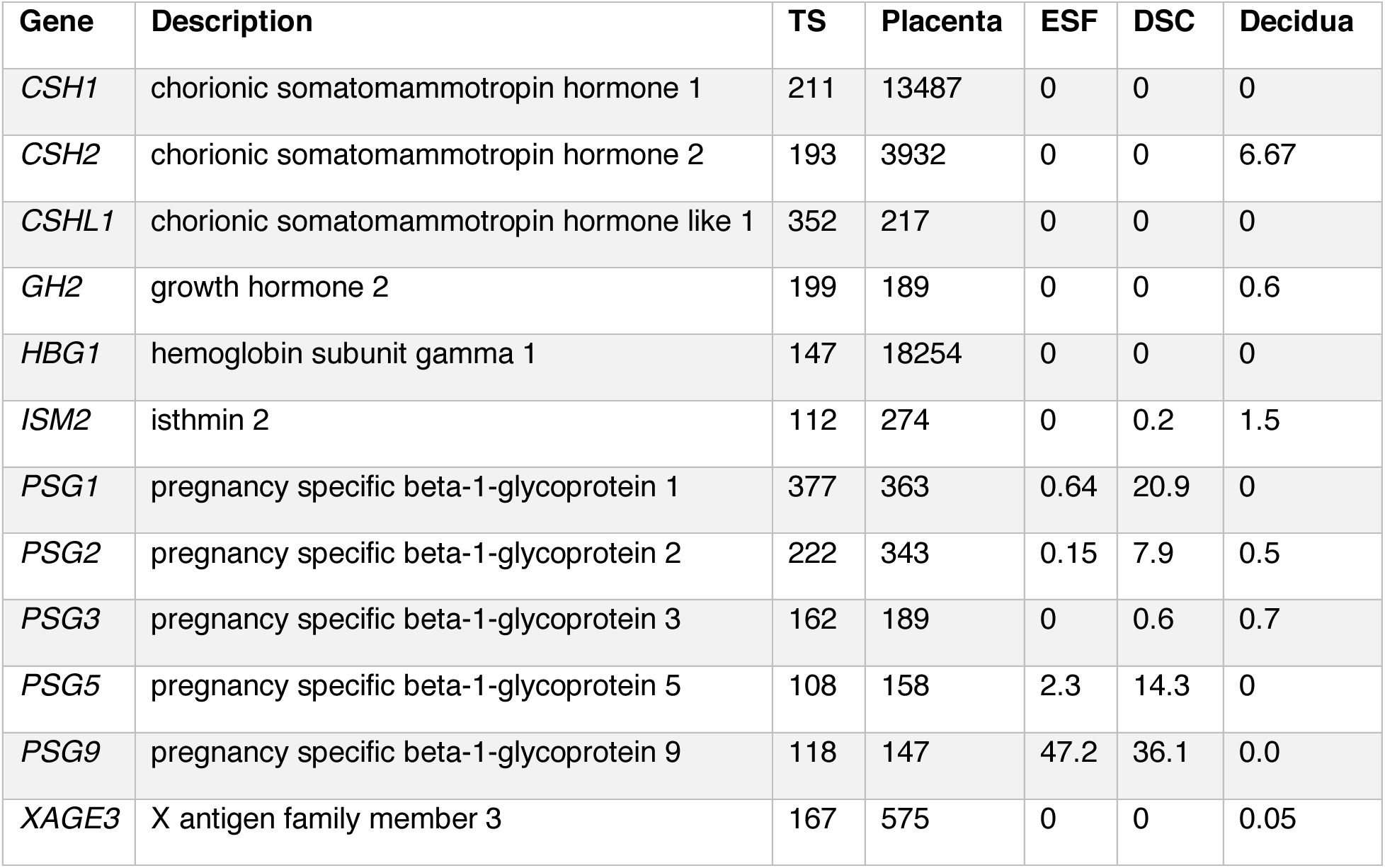
Expression of placental enriched genes in RNA-Seq data from human placenta, endometrial stromal fibroblasts (ESFs), decidual stromal cells (DSCs), and human first trimester decidua. Expression levels are shown as transcripts per million (TPM) values, the Tissue Specificity score (TS) is calculated as the fold enrichment of each gene relative to the tissue with the second highest expression of that gene. Placental data is from: https://www.proteinatlas.org/humanproteome/tissue/placenta.

| Gene | Description | TS | Placenta | ESF | DSC | Decidua |
| --- | --- | --- | --- | --- | --- | --- |
| *CSH1* | chorionic somatomammotropin hormone 1 | 211 | 13487 | 0 | 0 | 0 |
| *CSH2* | chorionic somatomammotropin hormone 2 | 193 | 3932 | 0 | 0 | 6.67 |
| *CSHL1* | chorionic somatomammotropin hormone like 1 | 352 | 217 | 0 | 0 | 0 |
| *GH2* | growth hormone 2 | 199 | 189 | 0 | 0 | 0.6 |
| *HBG1* | hemoglobin subunit gamma 1 | 147 | 18254 | 0 | 0 | 0 |
| *ISM2* | isthmin 2 | 112 | 274 | 0 | 0.2 | 1.5 |
| *PSG1* | pregnancy specific beta-1-glycoprotein 1 | 377 | 363 | 0.64 | 20.9 | 0 |
| *PSG2* | pregnancy specific beta-1-glycoprotein 2 | 222 | 343 | 0.15 | 7.9 | 0.5 |
| *PSG3* | pregnancy specific beta-1-glycoprotein 3 | 162 | 189 | 0 | 0.6 | 0.7 |
| *PSG5* | pregnancy specific beta-1-glycoprotein 5 | 108 | 158 | 2.3 | 14.3 | 0 |
| *PSG9* | pregnancy specific beta-1-glycoprotein 9 | 118 | 147 | 47.2 | 36.1 | 0.0 |
| *XAGE3* | X antigen family member 3 | 167 | 575 | 0 | 0 | 0.05 |

Next, we compared two different gene expression metrics to reconstruct the evolutionary history of endometrial gene expression: 1) Transcripts per million (TPM), a quantitative measure of gene expression that reflects the relative molar ratio of each transcripts in the transcriptome; and 2) Binary encoding, a discrete categorization of gene expression that classifies genes as expressed (state 1) or not expressed (state 0). For binary encoding we transformed transcript abundance estimates into discrete character states, such that genes with TPM ≥ 2.0 were coded as expressed (state 1), genes with TPM < 2.0 were coded as not expressed (state 0), and genes without data in specific species coded as missing (state ?); see Box 1 for a detailed justification of the TPM ≥ 2 cutoff. The TPM coded dataset grouped species randomly (Figure 1 – Figure supplement 1A), whereas the binary encoded endometrial gene expression dataset generally grouped species by phylogenetic relatedness (Figure 1 – Figure supplement 1B), suggesting greater signal to noise ratio than raw transcript abundance estimates. Therefore, we used the binary encoded endometrial transcriptome dataset to reconstruct ancestral gene expression states and trace the evolution of endometrial gene expression changes across vertebrate phylogeny (Figure 1A).

### Ancestral Transcriptome Reconstruction

Ancestral states for each gene were inferred with the empirical Bayesian method implemented in IQ-TREE 2 (Minh et al., 2020; Nguyen et al., 2015)using the species phylogeny shown in Figure 1A and the best-fitting model of character evolution determined by ModelFinder (Kalyaanamoorthy et al., 2017). The best fitting model was inferred to be the General Time Reversible model for binary data (GTR2), with character state frequencies optimized by maximum-likelihood (FO), and a FreeRate model of among site rate heterogeneity with four categories (R4) (Soubrier et al., 2012). We used ancestral transcriptome reconstructions to trace the evolution of gene expression gains (0 à 1) and losses (1 à 0) from the last common ancestor of mammals through to the Hominoid stem-lineage limiting our inferences to reconstructions with Bayesian posterior probabilities (BPP) ≥ 0.80 (Figure 1A and **Figure 1 – Source data 2**). Ancestral reconstructions with BPP ≥ 0.80 were excluded from over representation analyses.

### Data Exploration and Multi-Dimensional Scaling

We used classical Multi-Dimensional Scaling (MDS) to explore the structure of extant and ancestral transcriptomes. MDS is a multivariate data analysis method that can be used to visualize the similarity/dissimilarity between samples by plotting data points (in this case transcriptomes) onto two-dimensional plots. MDS returns an optimal solution that represents the data in a two-dimensional space, with the number of dimensions (k) specified *a priori*. Classical MDS preserves the original distance metric, between data points, as well as possible. MDS was performed using the vegan R package (Oksanen et al., 2019) with four reduced dimensions. Transcriptomes were grouped using K-means clustering with K=2-6, K=5 optimized the number of distinct clusters and cluster memberships (i.e., correctly grouping species by phylogenetic relationship, parity mode, and placenta type).

### scRNA-Seq Analyses

Maternal-fetus interface 10X Genomics scRNA-seq data was retrieved from the E-MTAB-6701 entry as a processed data matrix (Vento-Tormo et al., 2018). The RNA counts and major cell-type annotations were used as provided by the original publications. Seurat (v3.1.1), implemented in R (v3.6.0), was used for filtering, normalization, and cell-types clustering. The subclusters of cell types were annotated based on the known transcriptional markers from the literature survey. Briefly, we performed the following data processing steps: (1) Cells were filtered based on the criteria that individual cells must be expressing at least 1,000 and not more than 5,000 genes with a count ≥1; (2) Cells were filtered out if more than 5% of counts mapping to mitochondrial genes; (3) Data normalization was performed by dividing uniquely mapping read counts (defined by Seurat as unique molecular identified (UMI)) for each gene by the total number of counts in each cell and multiplying by 10,000. These normalized values were then log-transformed. Cell types were clustered by using the top 2000 variable genes expressed across all samples. Clustering was performed using the “FindClusters” function with essentially default parameters, except resolution was set to 0.1. The first 20 PCA dimensions were used in the construction of the shared-nearest neighbor (SNN) graph and the generation of 2-dimensional embeddings for data visualization using UMAP. Major cell types were assigned based on the original publication samples’ annotations, and cell sub-types within major cell types were annotated using the sub-cluster markers obtained from the above parameters. We then chose the decidual and PV cells to perform the single-cell trajectory, pseudotime analysis and cell ordering along an artificial temporal continuum analysis using Monocle2 (Qiu et al., 2017). The top 500 differentially expressed genes were used to distinguish between the sub-clusters of decidua and PV cell populations on pseudotime trajectory. The transcriptome from every single cell represents a pseudo-time point along an artificial time vector that denotes decidual and PV lineages’ progression, respectively. To compare the differentially expressed genes between *HTR2B*-positive and *HTR2B*-negative cells, we first divided the decidual and PV datasets into those groups of cells that either express *HTR2B* with a count ≥1 and those with zero counts. We then performed differentially expressed genes analysis between the mentioned two groups of cells using the bimodal test for significance.

To calculate the enrichment score of human-gain genes in each cell-types, we first transformed the data into a pseudobulk expression matrix by averaging all genes’ expression in each cell type. We then calculated the fraction of human-gained genes expressed (Observed) and the proportion of the rest of the genes expressed in each cell type (Expected). The enrichment ratio shown on the plot is the ratio of Observed and Expected values for each cell type. The P-value was calculated using a two-way Fisher exact test followed by Bonferroni correction. Specific codes to generate the data/plots are available upon request until we update it on GitHub.

### Over Representation Analyses (ORA)

We used WebGestalt v. 2019 (Liao et al., 2019) to identify enriched ontology terms using over-representation analysis (ORA). We used ORA to identify enriched terms for three pathway databases (KEGG, Reactome, and Wikipathway), three disease databases (Disgenet, OMIM, and GLAD4U), and a custom database of genes implicated in preterm birth by GWAS. The preterm birth gene set was assembled from the NHGRI-EBI Catalog of published genome-wide association studies (GWAS Catalog), including genes implicated in GWAS with either the ontology terms “Preterm Birth” (EFO_0003917) or “Spontaneous Preterm Birth” (EFO_0006917), as well as two recent preterm birth GWAS (Warrington *et al*., 2019; Sakabe *et al*., 2020) using a genome-wide significant P-value of 9×10^-6^. The custom gmt file used to test for enrichment of preterm birth associated genes is included as a supplementary data file to Figure 2 (**Figure 2 — Source data 1)**.

### Functional genomic analyses of the *HTR2B*, *PDCD1LG2*, and *CORIN* loci

We used previously published ChIP-Seq data generated from human DSCs that were downloaded from NCBI SRA and processed remotely using Galaxy (Afgan et al., 2016). ChIP-Seq reads were mapped to the human genome (GRCh37/hg19) using HISAT2 (Kim et al., 2019, 2015; Pertea et al., 2016)with default parameters and peaks called with MACS2 (Feng et al., 2012; Zhang et al., 2008) with default parameters. Samples included PLZF (GSE112362), H3K4me3 (GSE61793), H3K27ac (GSE61793), H3K4me1 (GSE57007), PGR (GSE94038), the PGR A and B isoforms (GSE62475), NR2F2 (GSE52008), FOSL2 (GSE94038), FOXO1 (GSE94037), PolII (GSE94037), GATA2 (GSE108408), SRC-2/NCOA2 (GSE123246), AHR (GSE114552), ATAC-Seq (GSE104720) and DNase1-Seq (GSE61793). FAIRE-Seq peaks were downloaded from the UCSC genome browser and not re-called.

We also used previously published RNA-Seq and microarray gene expression data generated from human ESFs and DSCs that were downloaded from NCBI SRA and processed remotely using Galaxy platform (https://usegalaxy.org/; Version 20.01) for RNA-Seq data and GEO2R for microarray data. RNA-Seq datasets were transferred from SRA to Galaxy using the Download and Extract Reads in FASTA/Q format from NCBI SRA tool (version 2.10.4+galaxy1). We used HISAT2 (version 2.1.0+galaxy5) to align reads to the Human hg38 reference genome using single-or paired-end options depending on the dataset and unstranded reads, and report alignments tailored for transcript assemblers including StringTie. Transcripts were assembled and quantified using StringTie (v1.3.6)(Pertea et al., 2016, 2015), with reference file to guide assembly and the “reference transcripts only” option, and output count files for differential expression with DESeq2/edgeR/limma-voom. Differentially expressed genes were identified using DESeq2 (Love et al., 2014)(version 2.11.40.6+galaxy1). The reference file for StringTie guided assembly was wgEncodeGencodeBasicV33. GEO2R performs comparisons on original submitter-supplied processed data tables using the GEOquery and limma R packages from the Bioconductor project (https://bioconductor.org/). These datasets included gene expression profiles of primary human ESFs treated for 48 hr with control non-targeting, PGR-targeting (GSE94036), FOXO1-targeting (GSE94036) or NR2F2 (COUP-TFII)-targeting (GSE47052) siRNA prior to decidualization stimulus for 72 hr; transfection with GATA2-targeting siRNA was followed immediately by decidualization stimulus (GSE108407). Probes were 206638_at (*HTR2B*), 220049_s_at (*PDCD1LG2*), and 220356_at (*CORIN*) for GSE4888 (endometrial gene expression throughout menstrual cycle) and 206638_at (*HTR2B*), 220049_s_at (*PDCD1LG2*), and 220356_at (*CORIN*) for GSE5999 (gene expression in basal plate throughout gestation). Multispecies RNA-Seq analysis of ESFs and DSCs is from GSE67659.

To assess chromatin looping, we utilized a previously published H3K27ac HiChIP dataset from a normal hTERT-immortalized endometrial cell line (E6E7hTERT) and three endometrial cancer cell lines (ARK1, Ishikawa and JHUEM-14) (O’Mara et al., 2019).

### Cell Culture and Serotonin (5-HT) Treatment

Human hTERT-immortalized endometrial stromal fibroblasts (T-HESC; CRL-4003, ATCC) were grown in maintenance medium, consisting of Phenol Red-free DMEM (31053–028, Thermo Fisher Scientific), supplemented with 10% charcoal-stripped fetal bovine serum (CS-FBS; 12676029, Thermo Fisher Scientific), 1% L-glutamine (25030–081, Thermo Fisher Scientific), 1% sodium pyruvate (11360070, Thermo Fisher Scientific), and 1x insulin-transferrin-selenium (ITS; 41400045, Thermo Fisher Scientific).

A total of 10^4^ ESFs were plated per well of a 96-well plate, 18 hours later cells were transfected in Opti-MEM (31985070, Thermo Fisher Scientific) with 100 ng of luciferase reporter plasmid, 10 ng Renilla control plasmid, 0.25 μl of Lipofectamine LTX (15338100, Thermo Fisher Scientific) and 0.1 μl Plus Reagent as per manufecturer’s protocol per well; final volume per well was 100 μl. Luciferase reporter plasmids were synthesized (GenScript) by cloning the response elements from the pGL4.29[luc2P/CRE/Hygro], pGL4.44[luc2P/AP1-RE/Hygro], pGL4.33[luc2P/SRE/Hygro], and pGL4.34[luc2P/SRF-RE/Hygro] plasmids into pGL3-Basic[minP] luciferase reporter. Unlike the pGL4 series vectors (Promega) that are hormone responsive, pGL3-Basic[minP] luciferase reporter includes a minimal promoter but is not hormone responsive. Final pathway reporter plasmids are: CRE_pGL3-Basic[minP] (cAMP/PKA), AP1_pGL3-Basic[minP] (AP1), SRE_pGL3-Basic[minP] (MAPK/ERK) and SRF_RE_pGL3-Basic[minP] (serum response factor).

ESFs were incubated in the transfection mixture for 6 hr. Then, ESFs were washed with warm PBS and incubated in the maintenance medium overnight. The next day, the medium in half of the wells was exchanged for the differentiation medium consisting of DMEM with Phenol Red and GlutaMAX (10566-024, Thermo Fisher Scientific), supplemented with 2% fetal bovine serum (FBS; 26140-079, Thermo Fisher Scientific), 1% sodium pyruvate (11360070, Thermo Fisher Scientific), 1 μM medroxyprogesterone 17-acetate (MPA; M1629, Sigma Aldrich) and 0.5 mM 8-Bromoadenosine 3′,5′-cyclic monophosphate (8-Br-cAMP; B5386, Sigma Aldrich). After 48 hrs, serotonin (5-HT; H9523, Sigma Aldrich) was added to the wells with both maintenance and differentiation medium (for each in triplicates) in the following concentrations: 50 μM, 200 μM and 1mM; vehicle control (0 μM) was water. After 6 hrs of incubation, we used a Dual Luciferase Reporter Assay (Promega) to quantify luciferase and Renilla luminescence following the manufacturer’s Dual Luciferase Reporter Assay protocol

## Acknowledgements

This study was supported by a grant from the March of Dimes (March of Dimes Prematurity Research Center) and a Burroughs Welcome Fund Preterm Birth Initiative grant (1013760) to principal investigator VJL. MS was supported by a presidential postdoctoral fellowship from Cornell University. The funders had no role in study design, data collection and analysis, decision to publish, or preparation of the manuscript.

---

**Figure 1 – Figure supplement 1.**
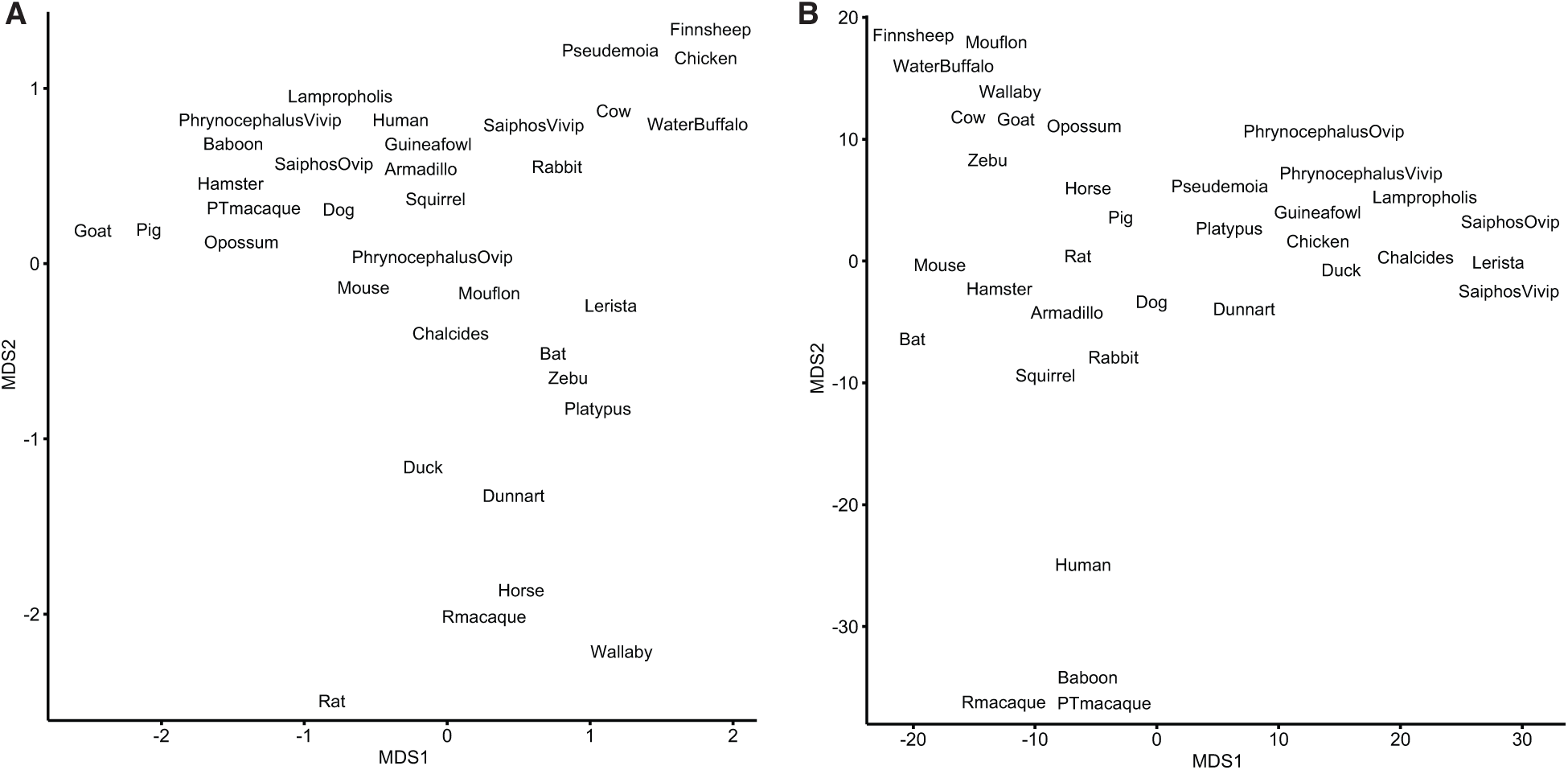
**(A)** Multidimensional scaling (MDS) plot of transcriptomes from extant species based on gene expression levels (TPMs) **(B)** Multidimensional scaling (MDS) plot of transcriptomes from extant species based on the binary encoded endometrial gene expression dataset. **Figure 1 — Source data 1. Species names (common and binomial), genome annotations used for RNA-Seq analysis, and parity mode.** **Figure 1 – Source data 2. Gene expression matrix and ancestral reconstruction results.**

**Figure 3 – Figure supplement 1.**
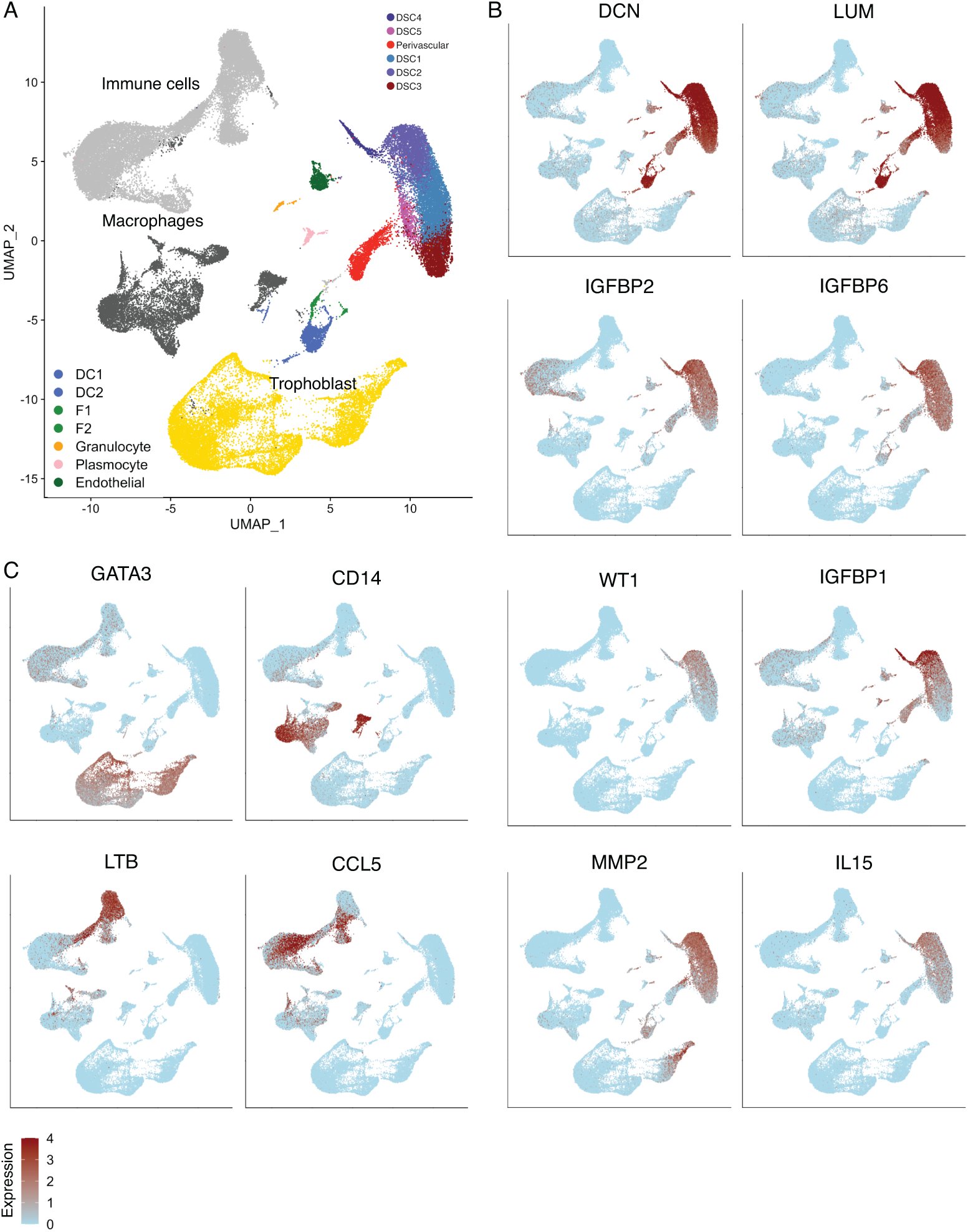
**(A)** UMAP clustering of cells from the first trimester maternal-fetal interface, major cell types and lineages are colored. **(B)** Feature plots based on the UMAP plot showing the single cell expression of marker genes for the endometrial stromal cell lineage. **(C)** Feature plots based on the UMAP plot showing the single cell expression of marker genes for immune cells (*LTB*, *CCL5*), macrophage (*CD14*), and trophoblasts (*GATA3*).

**Figure 3 – Figure supplement 2.**
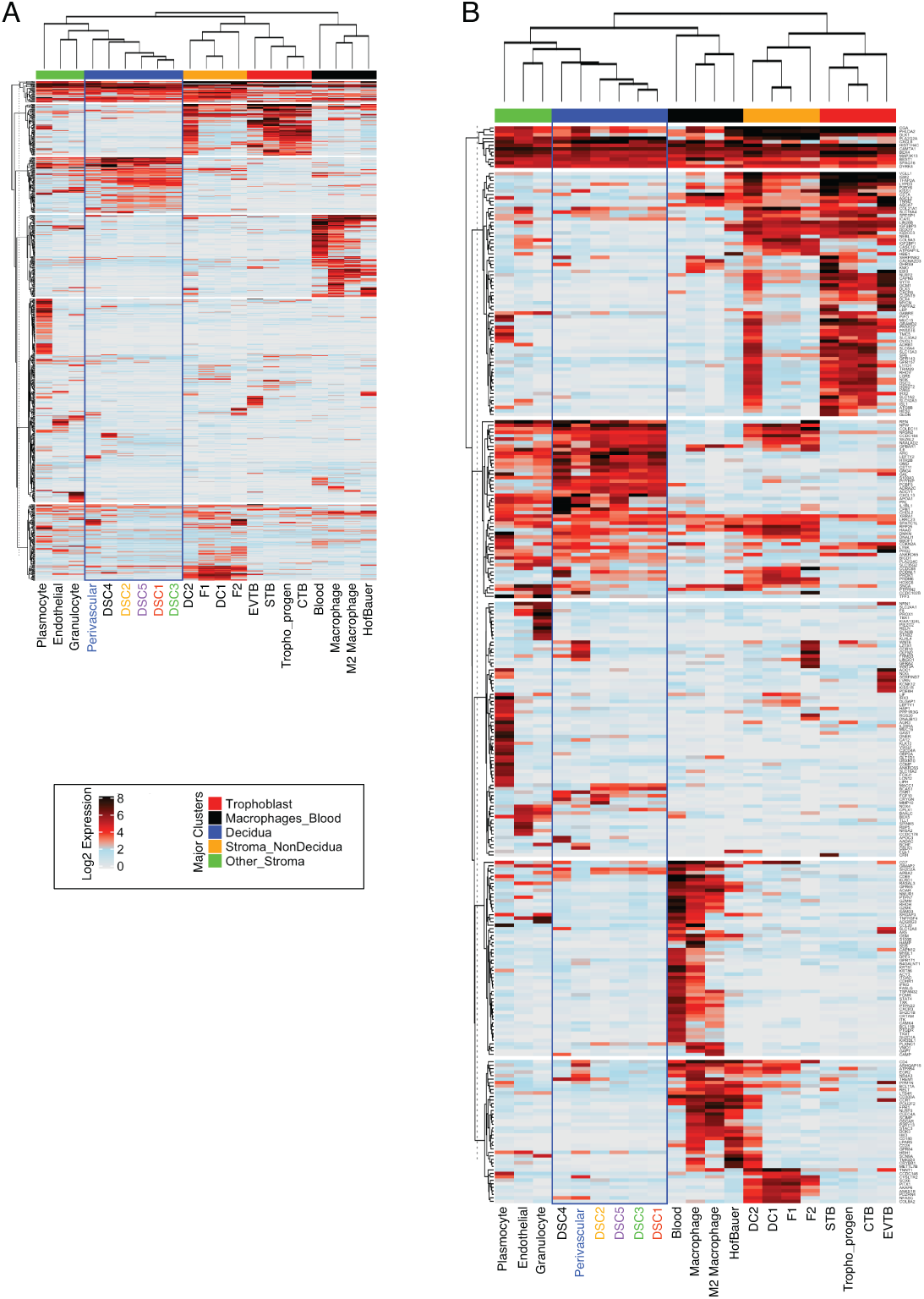
**(A)** Heatmap showing the expression of 840 human-gain genes in distinct cell-types identified as expressed at the maternal-fetus interface. Each row represents a gene that is expressed (log2 average scaled expression > 0) in at least one cell type denoted as individual columns. Prior to analysis, the single cellular expression of envelopes was averaged for each cell-types. The upper bars are manually annotated as clusters that denote the primary tissue where the cell-types are raised on the top of the heatmap. Rows are clustered using the k-means algorithm, and columns are hierarchically clustered using Euclidean distance and Spearman’s correlation. Endometrial stromal lineage cells are boxed in blue. **(B)** Heatmap similar to the one in A, except the analysis was performed on highly expressed 442 genes (Log2 scaled average expression > 0.5). Endometrial stromal lineage cells are boxed in blue.

**Figure 5 – Figure supplement 1.**
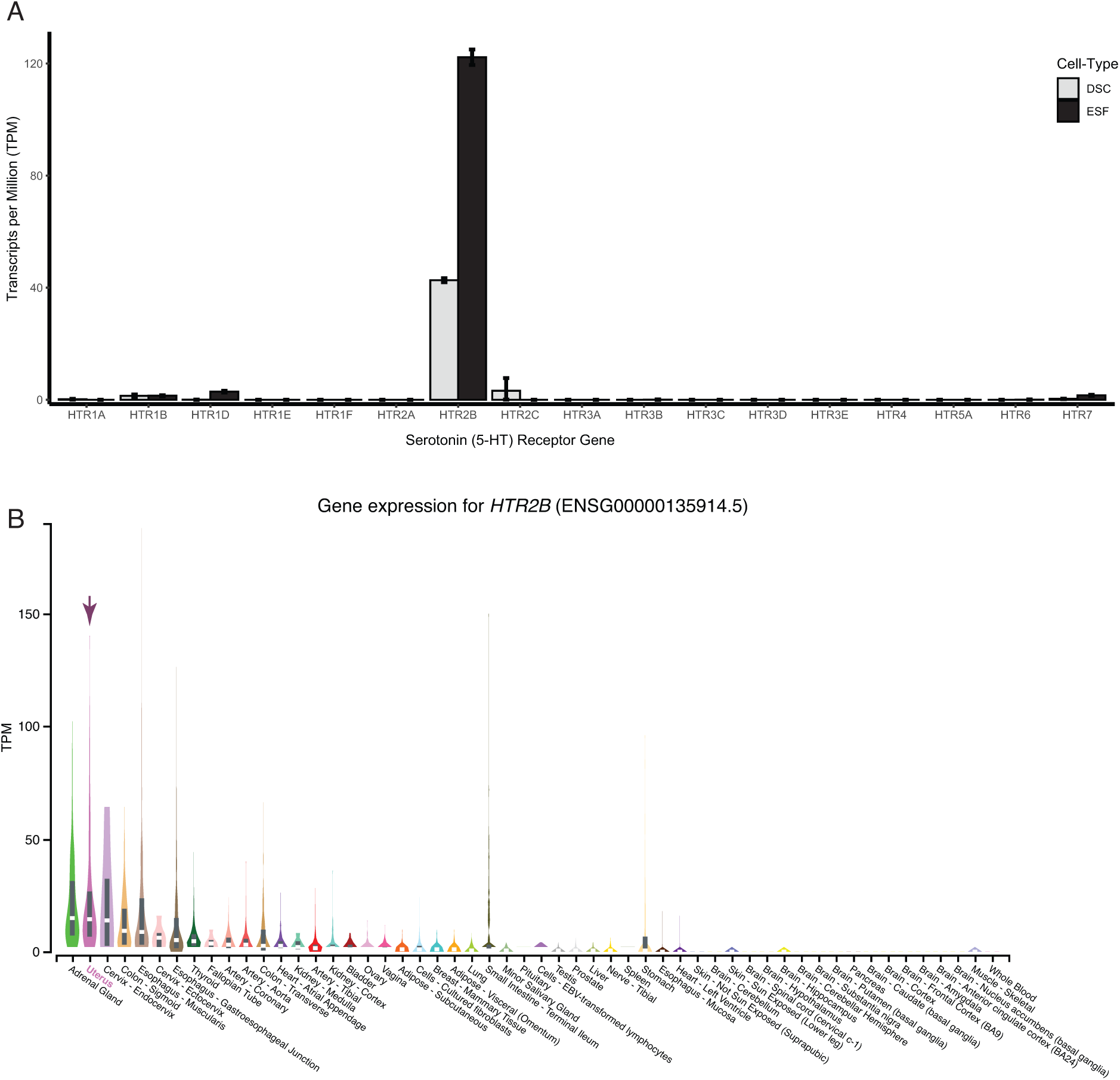
**(A)** *HTR2B* is the only serotonin (5-HT) receptor robustly expressed in human ESFs and DSCs. Expression of serotonin receptors in bulk RNA-Seq data (n = 2) from human endometrial stromal fibroblasts (ESF) and ESFs induced into decidual stromal cells (DSC) by cAMP/progesterone treatment for 48 hours. **(B)** Expression of *HTR2B* in GTEx tissues. Violin plots are colored by anatomical system; the uterus is indicated by an arrow. **Figure 5 – Source data 1.** Genes that are differentially expressed between *HTR2B*^+^ and *HTR2B*^−^ cells, and the pathways/disease ontologies in which they are enriched.

**Figure 6 – Figure supplement 1.**
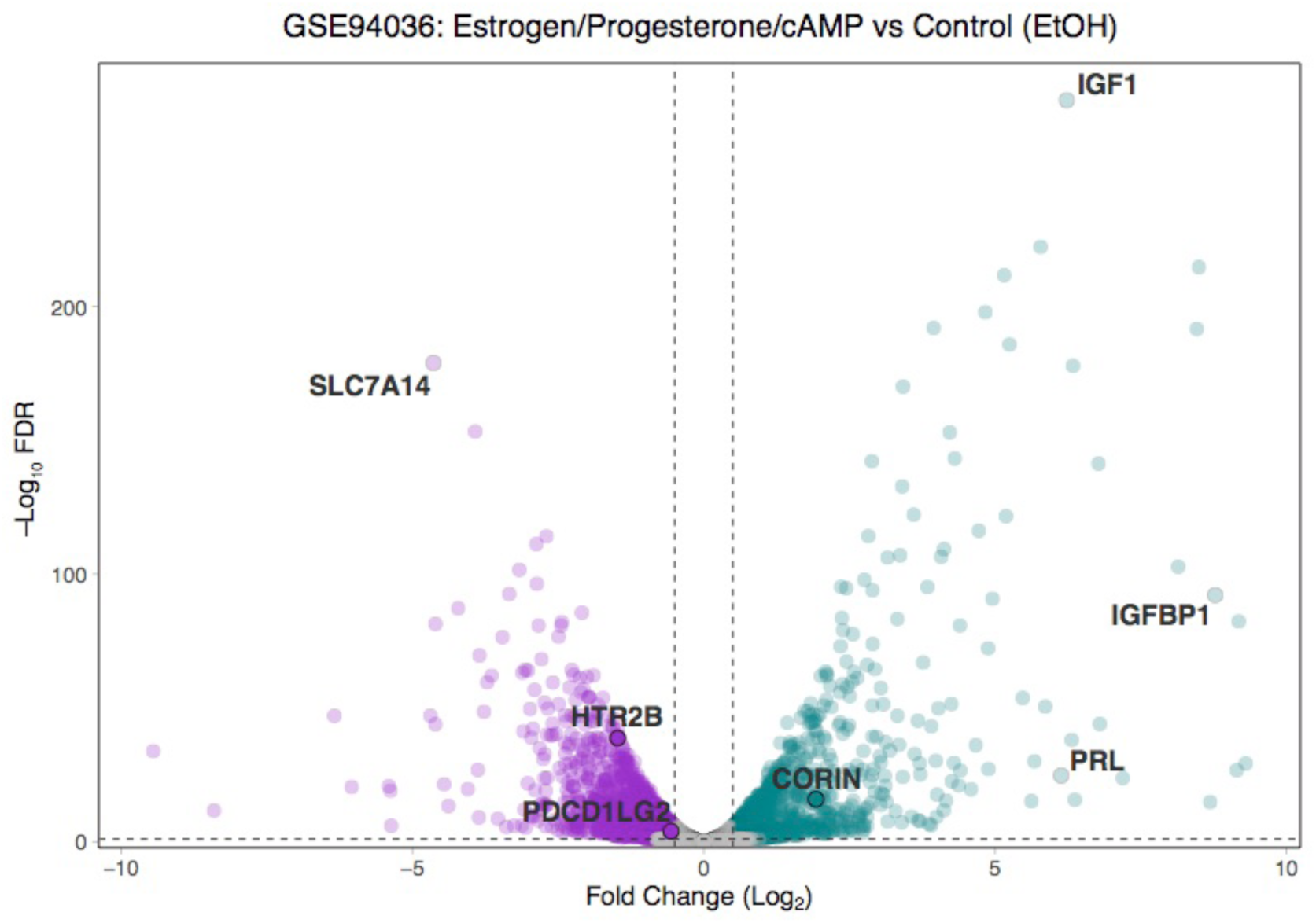
Volcano plot of gene expression changes between ESFs and DSCs. Differentially expressed genes at −Log_10_ FDR ≤ 2 and Log_2_ Fold-Change ≥ 0.5/-0.5 are colored purple (down-regulated) and green (up-regulated). *HTR2B*, *PDCD1LG2*, and *CORIN* are shown, as are classic marker genes of decidualization (*IGF1*, *IGFBP1*, and *PRL*).

**Figure 6 – Figure supplement 2.**
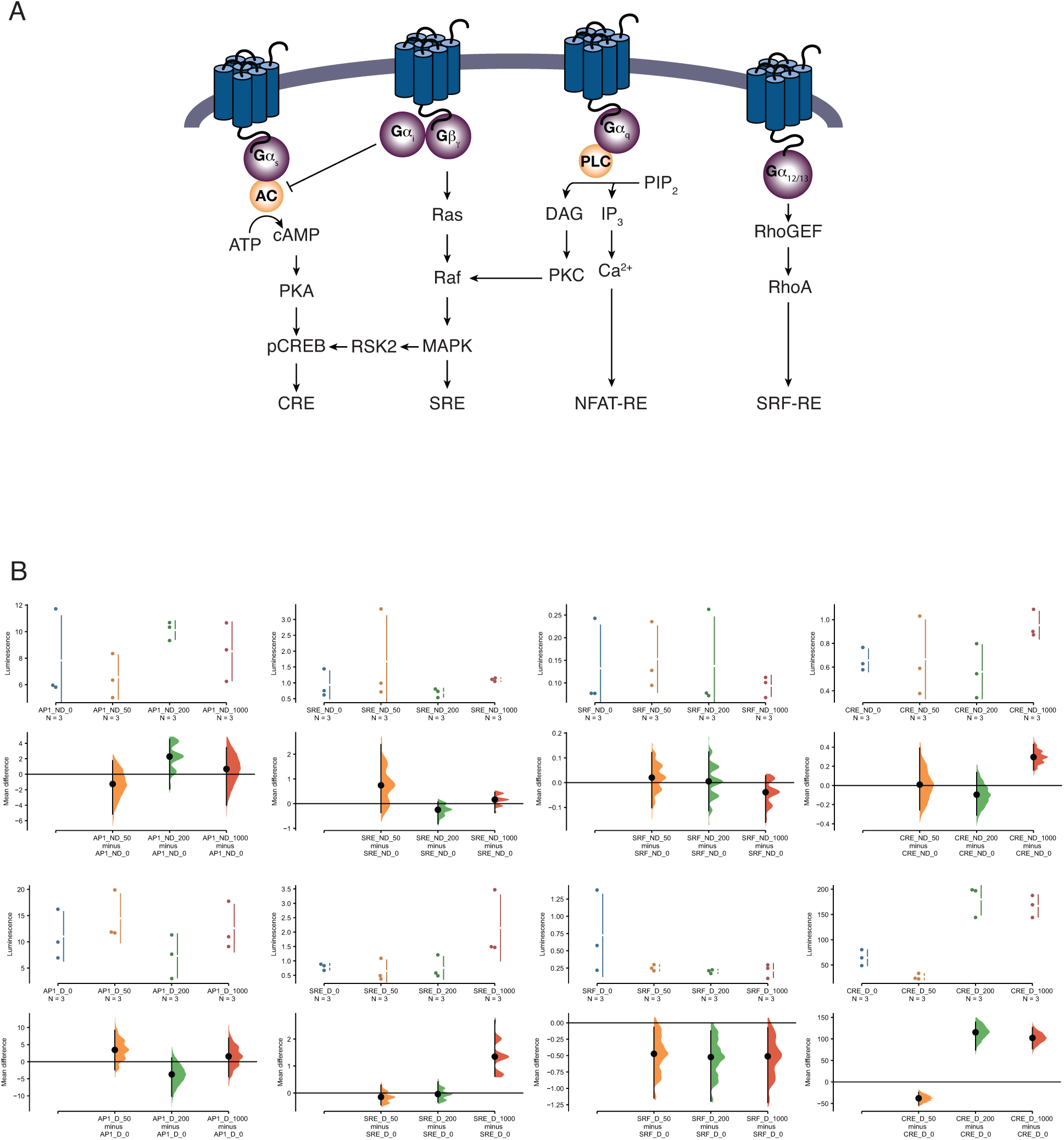
Pathway activation screen. **(A)** Schematic of the luciferase-based pathway reporter system. Pathways leading to luciferase expression from the cAMP/PKA (pGL4.29[luc2P/CRE/Hygro]), AP1 (pGL4.44[luc2P/AP1-RE/Hygro]), MAPK/ERK (pGL4.33[luc2P/SRE/Hygro]), and Serum Response Factor Response Element (pGL4.34[luc2P/SRF-RE/Hygro]) reporter vectors upon serotonin treatment are shown. **(B)** Cumming estimation plots showing mean difference in luminescence for the serotonin dose response from pathway reporter vectors. Upper plots show relative luminescence of human ESFs (ND for non-decidualized) or DSCs (D for decidualized) transiently transfected with a luciferase expression vector that drives the transcription of the indicated luciferase reporter gene 6 hours after treatment with serotonin (50, 200 and 1000 μM) or vehicle control (0). Lower plots, mean differences are plotted as bootstrap sampling distributions (n = 5000; the confidence interval is bias-corrected and accelerated). Each mean difference is depicted as a dot. Each 95% confidence interval is indicated by the vertical error bars. P-values indicate the likelihoods of observing the effect sizes, if the null hypothesis of zero difference is true. AP1 ESF Results: The unpaired mean difference between AP1_ND_0 and AP1_ND_50 is -1.26 [95.0%CI - 5.14, 1.77]. The P value of the two-sided permutation t-test is 0.652. The unpaired mean difference between AP1_ND_0 and AP1_ND_200 is 2.28 [95.0%CI - 1.93, 4.54]. The P value of the two-sided permutation t-test is 0.292. The unpaired mean difference between AP1_ND_0 and AP1_ND_1000 is 0.684 [95.0%CI -3.99, 3.44]. The P value of the two-sided permutation t-test is 0.693. AP1 DSC Results: The unpaired mean difference between AP1_D_0 and AP1_D_50 is 3.45 [95.0%CI -2.37, 9.23]. The P value of the two-sided permutation t-test is 0.31. The unpaired mean difference between AP1_D_0 and AP1_D_200 is -3.72 [95.0%CI - 10.1, 1.14]. The P value of the two-sided permutation t-test is 0.392. The unpaired mean difference between AP1_D_0 and AP1_D_1000 is 1.56 [95.0%CI - 4.41, 6.91]. The P value of the two-sided permutation t-test is 0.528. SRF ESF Results: The unpaired mean difference between SRF_ND_0 and SRF_ND_50 is 0.0203 [95.0%CI -0.101, 0.122]. The P value of the two-sided permutation t-test is 0.516. The unpaired mean difference between SRF_ND_0 and SRF_ND_200 is 0.00516 [95.0%CI -0.114, 0.124]. The P value of the two-sided permutation t-test is 0.727. The unpaired mean difference between SRF_ND_0 and SRF_ND_1000 is -0.0387 [95.0%CI -0.16, 0.0276]. The P value of the two-sided permutation t-test is 0.693. SRF DSC Results: The unpaired mean difference between SRF_D_0 and SRF_D_50 is -0.473 [95.0%CI - 1.15, -0.0678]. The P value of the two-sided permutation t-test is 0.2. The unpaired mean difference between SRF_D_0 and SRF_D_200 is -0.523 [95.0%CI - 1.19, -0.122]. The P value of the two-sided permutation t-test is 0.104. The unpaired mean difference between SRF_D_0 and SRF_D_1000 is -0.511 [95.0%CI - 1.22, -0.0742]. The P value of the two-sided permutation t-test is 0.199. SRE ESF Results: The unpaired mean difference between SRE_ND_0 and SRE_ND_50 is 0.74 [95.0%CI - 0.361, 2.39]. The P value of the two-sided permutation t-test is 0.494. The unpaired mean difference between SRE_ND_0 and SRE_ND_200 is -0.248 [95.0%CI -0.816, 0.0713]. The P value of the two-sided permutation t-test is 0.403. The unpaired mean difference between SRE_ND_0 and SRE_ND_1000 is 0.164 [95.0%CI -0.361, 0.459]. The P value of the two-sided permutation t-test is 0.405. SRE DSC Results: The unpaired mean difference between SRE_D_0 and SRE_D_50 is -0.147 [95.0%CI - 0.44, 0.293]. The P value of the two-sided permutation t-test is 0.501. The unpaired mean difference between SRE_D_0 and SRE_D_200 is -0.0399 [95.0%CI -0.336, 0.412]. The P value of the two-sided permutation t-test is 0.793. The unpaired mean difference between SRE_D_0 and SRE_D_1000 is 1.35 [95.0%CI 0.624, 2.69]. The P value of the two-sided permutation t-test is 0.0. CRE ESF Results: The unpaired mean difference between CRE_ND_0 and CRE_ND_50 is 0.00845 [95.0%CI -0.255, 0.39]. The P value of the two-sided permutation t-test is 0.798. The unpaired mean difference between CRE_ND_0 and CRE_ND_200 is -0.096 [95.0%CI -0.31, 0.135]. The P value of the two-sided permutation t-test is 0.41. The unpaired mean difference between CRE_ND_0 and CRE_ND_1000 is 0.296 [95.0%CI 0.161, 0.43]. The P value of the two-sided permutation t-test is 0.0. CRE DSC Results: The unpaired mean difference between CRE_D_0 and CRE_D_50 is -37.8 [95.0%CI - 53.7, -24.1]. The P value of the two-sided permutation t-test is 0.0. The unpaired mean difference between CRE_D_0 and CRE_D_200 is 1.15e+02 [95.0%CI 74.2, 1.39e+02]. The P value of the two-sided permutation t-test is 0.0. The unpaired mean difference between CRE_D_0 and CRE_D_1000 is 1.02e+02 [95.0%CI 77.3, 1.26e+02]. The P value of the two-sided permutation t-test is 0.0.

**Figure 7 – Figure supplement 1.**
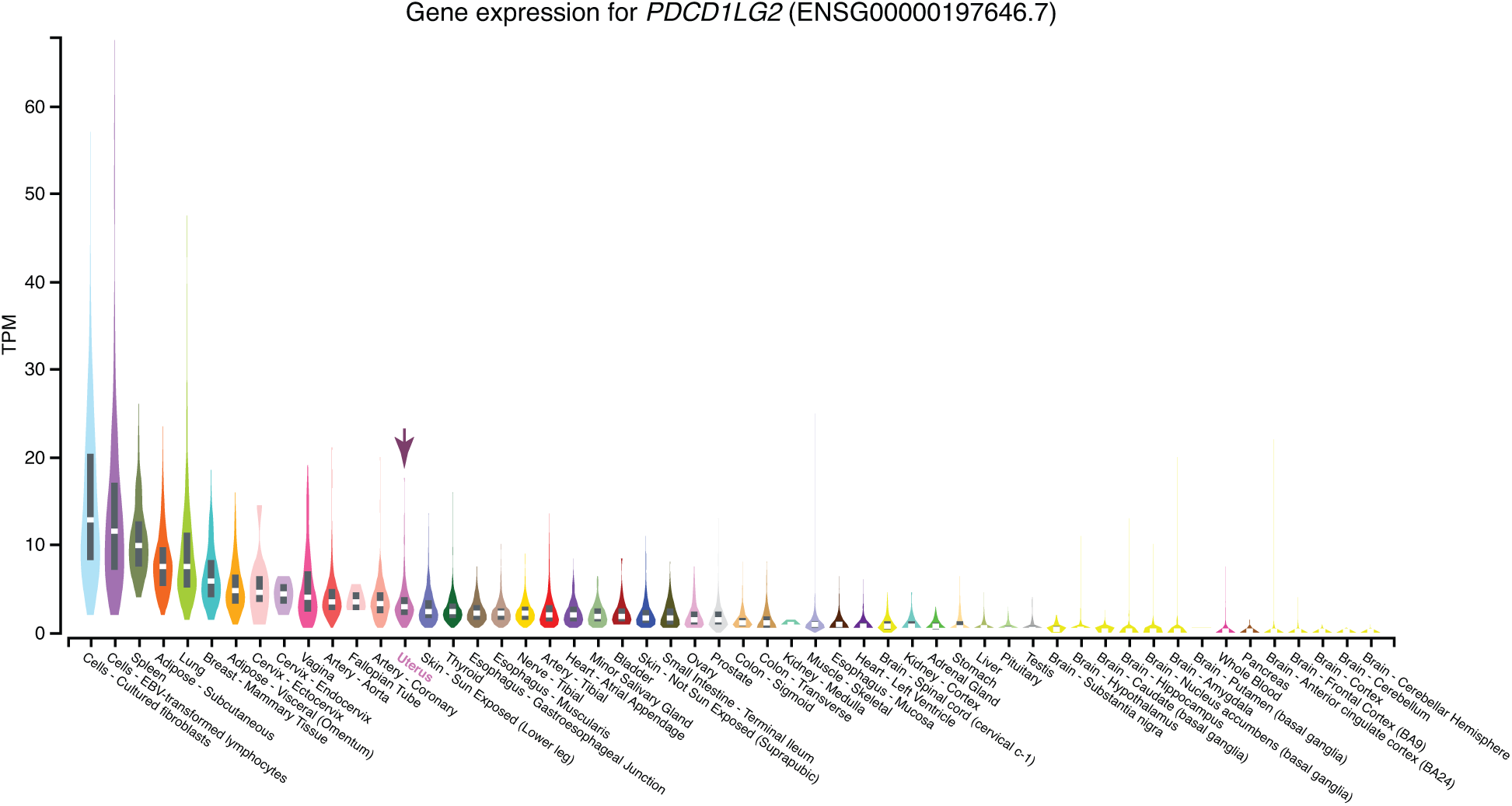
Expression of *PDCD1LG2* in GTEx tissues. Violin plots are colored by anatomical system; the uterus is indicated by an arrow.

**Figure 8 – Figure supplement 1.**
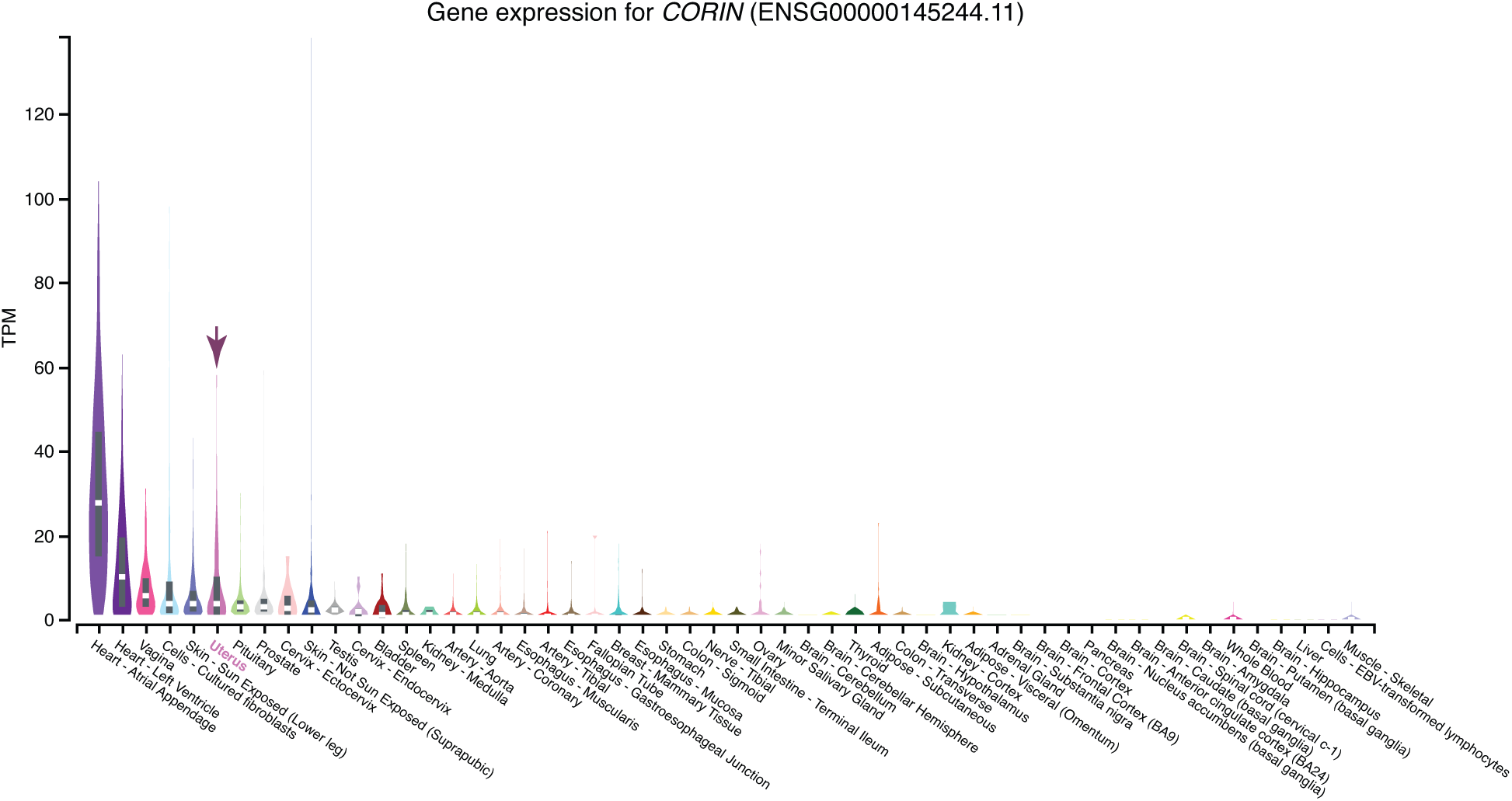
Expression of *CORIN* in GTEx tissues. Violin plots are colored by anatomical system; the uterus is indicated by an arrow.

## References

Abbas Y, Brunel LG, Hollinshead MS, Fernando RC, Gardner L, Duncan I, Moffett A, Best S, Turco MY, Burton GJ, Cameron RE. 2020. Generation of a three-dimensional collagen scaffold-based model of the human endometrium. Interface Focus 10:20190079. doi:10.1098/rsfs.2019.0079

Abbot P, Rokas A. 2017. Mammalian pregnancy. Curr Biol 27:R127–R128. doi:10.1016/j.cub.2016.10.046

Afgan E, Baker D, Beek M van den, Blankenberg D, Bouvier D, Čech M, Chilton J, Clements D, Coraor N, Eberhard C, Grüning B, Guerler A, Hillman-Jackson J, Kuster GV, Rasche E, Soranzo N, Turaga N, Taylor J, Nekrutenko A, Goecks J. 2016. The Galaxy platform for accessible, reproducible and collaborative biomedical analyses: 2016 update. Nucleic Acids Res 44:W3–W10. doi:10.1093/nar/gkw343

Armstrong DL, McGowen MR, Weckle A, Pantham P, Caravas J, Agnew D, Benirschke K, Savage-Rumbaugh S, Nevo E, Kim CJ, Wagner GP, Romero R, Wildman DE. 2017. The core transcriptome of mammalian placentas and the divergence of expression with placental shape. Placenta 57:71–78. doi:10.1016/j.placenta.2017.04.015

Aschebrook-Kilfoy B, Argos M, Pierce BL, Tong L, Jasmine F, Roy S, Parvez F, Ahmed A, Islam T, Kibriya MG, Ahsan H. 2015. Genome-wide association study of parity in Bangladeshi women. Plos One 10:e0118488. doi:10.1371/journal.pone.0118488

Behringer RR, Eakin GS, Renfree MB. 2006. Mammalian diversity: gametes, embryos and reproduction. Reproduction, fertility, and development 18:99–107.

Benton ML, Abraham A, LaBella AL, Abbot P, Rokas A, Capra JA. 2021. The influence of evolutionary history on human health and disease. Nat Rev Genet 1–15. doi:10.1038/s41576-020-00305-9

Boretto M, Cox B, Noben M, Hendriks N, Fassbender A, Roose H, Amant F, Timmerman D, Tomassetti C, Vanhie A, Meuleman C, Ferrante M, Vankelecom H. 2017. Development of organoids from mouse and human endometrium showing endometrial epithelium physiology and long-term expandability. Development 144:1775–1786. doi:10.1242/dev.148478

Bourne GH. 1970. The Chimpanzee: Immunology, infections, hormones, anatomy, and behavior of chimpanzees. University Park Press.

Bray NL, Pimentel H, Melsted P, Pachter L. 2016. Near-optimal probabilistic RNA-seq quantification. Nat Biotechnol 34:525–527. doi:10.1038/nbt.3519

Burley N. 1979. The Evolution of Concealed Ovulation. The American Naturalist 114:835–858.

Carter AM. 2011. Comparative studies of placentation and immunology in non-human primates suggest a scenario for the evolution of deep trophoblast invasion and an explanation for human pregnancy disorders. Reproduction 141:391–396. doi:10.1530/rep-10-0530

Carter AM, Enders AC, Pijnenborg R. 2015. The role of invasive trophoblast in implantation and placentation of primates. Philosophical Transactions Royal Soc B Biological Sci 370:20140070. doi:10.1098/rstb.2014.0070

Carter AM, Mess AM. 2017. The evolution of fetal membranes and placentation in carnivores and ungulates (Ferungulata). Anim Reprod 14:124–135. doi:10.21451/1984-3143-ar903

Chen Y, Palm F, Lesch K-P, Gerlach M, Moessner R, Sommer C. 2011. 5-hydroxyindolacetic acid (5-HIAA), a main metabolite of serotonin, is responsible for complete Freund’s adjuvant-induced thermal hyperalgesia in mice. Mol Pain 7:21. doi:10.1186/1744-8069-7-21

Clark KE, Mills EG, Otte TE, Stys SJ. 1980. Effect of serotonin on uterine blood flow in pregnant and nonpregnant sheep. Life Sciences 27:2655–2661.

Crosley EJ, Elliot MG, Christians JK, Crespi BJ. 2013. Placental invasion, preeclampsia risk and adaptive molecular evolution at the origin of the great apes: evidence from genome-wide analyses. Placenta 34:127–132. doi:10.1016/j.placenta.2012.12.001

Csapo AI. 1956. Progesterone block. Am J Anat 98:273–291. doi:10.1002/aja.1000980206

Csapo AI, Pinto-Dantas P-DCA. 1965. The effect of progesterone on the human uterus. Proc National Acad Sci 54:1069–1076. doi:10.1073/pnas.54.4.1069

Cui Y, Wang W, Dong N, Lou J, Srinivasan DK, Cheng W, Huang X, Liu M, Fang C, Peng J, Chen S, Wu S, Liu Z, Dong L, Zhou Y, Wu Q. 2012. Role of corin in trophoblast invasion and uterine spiral artery remodelling in pregnancy. Nature 484:246–250. doi:10.1038/nature10897

Cummins J. 1999. Evolutionary forces behind human infertility. Nature 397:557–558. doi:10.1038/17471

Eke A, Saccone G, Berghella V. 2016. Selective serotonin reuptake inhibitor (SSRI) use during pregnancy and risk of preterm birth: a systematic review and meta-analysis. Bjog Int J Obstetrics Gynaecol 123:1900–1907. doi:10.1111/1471-0528.14144

Elliot MG. 2017. Evolutionary origins of preeclampsia. Pregnancy Hypertens Int J Women’s Cardiovasc Heal 7:56. doi:10.1016/j.preghy.2016.10.006

Emera D, Casola C, Lynch VJ, Wildman DE, Agnew D, Wagner GP. 2012a. Convergent Evolution of Endometrial Prolactin Expression in Primates, Mice, and Elephants Through the Independent Recruitment of Transposable Elements. Mol Biol Evol 29:239–247. doi:10.1093/molbev/msr189

Emera D, Romero R, Wagner G. 2012b. The evolution of menstruation: A new model for genetic assimilation. Bioessays 34:26–35. doi:10.1002/bies.201100099

Feng J, Liu T, Qin B, Zhang Y, Liu XS. 2012. Identifying ChIP-seq enrichment using MACS. Nat Protoc 7:1728–1740. doi:10.1038/nprot.2012.101

Finn CA. 1998. Menstruation: A Nonadaptive Consequence of Uterine Evolution. Q Rev Biology 73:163–173. doi:10.1086/420183

Freyer C, Renfree MB. 2009. The mammalian yolk sac placenta. J Exp Zoology Part B Mol Dev Evol 312B:545–554. doi:10.1002/jez.b.21239

Freyer C, Zeller U, Renfree MB. 2003. The marsupial placenta: A phylogenetic analysis. J Exp Zool 299A:59–77. doi:10.1002/jez.a.10291

Gellersen B, Brosens IA, Brosens JJ. 2007. Decidualization of the human endometrium: mechanisms, functions, and clinical perspectives. Semin Reprod Med 25:445–453. doi:10.1055/s-2007-991042

Gellersen B, Brosens J. 2003. Cyclic AMP and progesterone receptor cross-talk in endometrium: a decidualizing affair. J Endocrinology 178:357–372.

Gerlo S, Davis JRE, Mager DL, Kooijman R. 2006. Prolactin in man: a tale of two promoters. Bioessays 28:1051–1055. doi:10.1002/bies.20468

Ghiotto M, Gauthier L, Serriari N, Pastor S, Truneh A, Nunès JA, Olive D. 2010. PD-L1 and PD-L2 differ in their molecular mechanisms of interaction with PD-1. Int Immunol 22:651–660. doi:10.1093/intimm/dxq049

Giresi PG, Kim J, McDaniell RM, Iyer VR, Lieb JD. 2007. FAIRE (Formaldehyde-Assisted Isolation of Regulatory Elements) isolates active regulatory elements from human chromatin. Genome Res 17:877–885. doi:10.1101/gr.5533506

Grzeskowiak LE, Gilbert AL, Morrison JL. 2012. Neonatal outcomes after late-gestation exposure to selective serotonin reuptake inhibitors. J Clin Psychopharm 32:615–621. doi:10.1097/jcp.0b013e31826686bc

Hebenstreit D, Fang M, Gu M, Charoensawan V, Oudenaarden A van, Teichmann SA. 2011. RNA sequencing reveals two major classes of gene expression levels in metazoan cells. Mol Syst Biol 7. doi:10.1038/msb.2011.28

Hou Z, Romero R, Uddin M, Than NG, Wildman DE. 2009. Adaptive history of single copy genes highly expressed in the term human placenta. Genomics 93:33–41. doi:10.1016/j.ygeno.2008.09.005

Hughes RL, Hall LS. 1998. Early development and embryology of the platypus. Philosophical Transactions Royal Soc Lond Ser B Biological Sci 353:1101–1114. doi:10.1098/rstb.1998.0269

Huybrechts KF, Sanghani RS, Avorn J, Urato AC. 2014. Preterm Birth and Antidepressant Medication Use during Pregnancy: A Systematic Review and Meta-Analysis. Plos One 9:e92778. doi:10.1371/journal.pone.0092778

Jones JB, Pycock CJ. 1978. Aminotic fluid levels of 5-hydroxytryptamine and 5-hydroxyndoleacetic acid before and during labour. Brit J Obstet Gynaec 85:530–32.

Kalyaanamoorthy S, Minh BQ, Wong TKF, Haeseler A von, Jermiin LS. 2017. ModelFinder: fast model selection for accurate phylogenetic estimates. Nat Methods 14:587–589. doi:10.1038/nmeth.4285

Keeling M, Roberts J. 1972. Histology, Reproduction, and Restraint.

Kim D, Langmead B, Salzberg SL. 2015. HISAT: a fast spliced aligner with low memory requirements. Nat Methods 12:357–360. doi:10.1038/nmeth.3317

Kim D, Paggi JM, Park C, Bennett C, Salzberg SL. 2019. Graph-based genome alignment and genotyping with HISAT2 and HISAT-genotype. Nat Biotechnol 37:907–915. doi:10.1038/s41587-019-0201-4

Kin K, Maziarz J, Chavan AR, Kamat M, Vasudevan S, Birt A, Emera D, Lynch VJ, Ott TL, Pavlicev M. 2016. The transcriptomic evolution of mammalian pregnancy: gene expression innovations in endometrial stromal fibroblasts. Genome biology and evolution 8:2459–2473.

Kin K, Nnamani MC, Lynch VJ, Michaelides E, Wagner GP. 2015. Cell-type phylogenetics and the origin of endometrial stromal cells. Cell reports 10:1398–1409.

Klein C, Roussel G, Brun S, Rusu C, Patte-Mensah C, Maitre M, Mensah-Nyagan A-G. 2018. 5-HIAA induces neprilysin to ameliorate pathophysiology and symptoms in a mouse model for Alzheimer’s disease. Acta Neuropathologica Commun 6:136. doi:10.1186/s40478-018-0640-z

Kliman HJ, Quaratella SB, Setaro AC, Siegman EC, Subha ZT, Tal R, Milano KM, Steck TL. 2018. Pathway of Maternal Serotonin to the Human Embryo and Fetus. Endocrinology 159:1609–1629. doi:10.1210/en.2017-03025

Kolstad KD, Mayo JA, Chung L, Chaichian Y, Kelly VM, Druzin M, Stevenson DK, Shaw GM, Simard JF. 2019. Preterm birth phenotypes in women with autoimmune rheumatic diseases: a population-based cohort study. Bjog Int J Obstetrics Gynaecol 31:9. doi:10.1111/1471-0528.15970

Koren Z, Eckstein B, Brzezinski A. 1961. Adrenaline, noradrenaline and serotonin estimations in urine and amniotic fluid during delivery. BJOG: An International ….

Koren Z, Pfeifer Y, Sulman FG. 1966. INDUCTION OF LEGAL ABORTION BY INTRA-UTERINE INSTILLATION OF PARGYLINE HYDROCHLORIDE (EUTONYL). Reproduction 12:75–79. doi:10.1530/jrf.0.0120075

Koren Z, Pfeifer Y, Sulman FG. 1965. Serotonin content of human placenta and fetus during pregnancy. Am J Obstet Gynecol 93:411–415. doi:10.1016/0002-9378(65)90070-0

Kosova G, Stephenson MD, Lynch VJ, Ober C. 2015. Evolutionary forward genomics reveals novel insights into the genes and pathways dysregulated in recurrent early pregnancy loss. Human Reproduction 30:519–529.

LaBella AL, Abraham A, Pichkar Y, Fong SL, Zhang G, Muglia LJ, Abbot P, Rokas A, Capra JA. 2020. Accounting for diverse evolutionary forces reveals mosaic patterns of selection on human preterm birth loci. Nat Commun 11:3731. doi:10.1038/s41467-020-17258-6

Latchman Y, Wood CR, Chernova T, Chaudhary D, Borde M, Chernova I, Iwai Y, Long AJ, Brown JA, Nunes R, Greenfield EA, Bourque K, Boussiotis VA, Carter LL, Carreno BM, Malenkovich N, Nishimura H, Okazaki T, Honjo T, Sharpe AH, Freeman GJ. 2001. PD-L2 is a second ligand for PD-1 and inhibits T cell activation. Nat Immunol 2:261–268. doi:10.1038/85330

Laurent L, Deroy K, St-Pierre J, Côté F, Sanderson JT, Vaillancourt C. 2017. Human placenta expresses both peripheral and neuronal isoform of tryptophan hydroxylase. Biochimie 140:159–165. doi:10.1016/j.biochi.2017.07.008

Liao Y, Wang J, Jaehnig EJ, Shi Z, Zhang B. 2019. WebGestalt 2019: gene set analysis toolkit with revamped UIs and APIs. Nucleic Acids Res 47:W199–W205. doi:10.1093/nar/gkz401

Loose R, Paterson WG. 1966. 5-Hydroxyindole Acetic Acid In Amniotic Fluid And Foetal 5-Hydroxytryptamine Metabolism. Bjog Int J Obstetrics Gynaecol 73:647–653. doi:10.1111/j.1471-0528.1966.tb15546.x

Love MI, Huber W, Anders S. 2014. Moderated estimation of fold change and dispersion for RNA-seq data with DESeq2. Genome Biol 15.

Lynch VJ, Nnamani MC, Kapusta A, Brayer K, Plaza SL, Mazur EC, Emera D, Sheikh SZ, Grutzner F, Bauersachs S, Graf A, Young SL, Lieb JD, DeMayo FJ, Feschotte C, Wagner GP. 2015. Ancient transposable elements transformed the uterine regulatory landscape and transcriptome during the evolution of mammalian pregnancy. Cell Reports 10:551–561. doi:10.1016/j.celrep.2014.12.052

Lynch VJ, Tanzer A, Wang Y, Leung FC, Gellersen B, Emera D, Wagner GP. 2008. Adaptive changes in the transcription factor HoxA-11 are essential for the evolution of pregnancy in mammals. Proc National Acad Sci 105:14928–14933. doi:10.1073/pnas.0802355105

Marinić M, Mika K, Chigurupati S, Lynch VJ. 2021. Evolutionary transcriptomics implicates HAND2 in the origins of implantation and regulation of gestation length. Elife 10:e61257. doi:10.7554/elife.61257

Marinić M, Rana S, Lynch VJ. 2020. Derivation of endometrial gland organoids from term placenta. Placenta 101:75–79. doi:10.1016/j.placenta.2020.08.017

Marshall SA, Hannan NJ, Jelinic M, Nguyen TPH, Girling JE, Parry LJ. 2018. Animal models of preeclampsia: translational failings and why. Am J Physiology-regulatory Integr Comp Physiology 314:R499–R508. doi:10.1152/ajpregu.00355.2017

Mazur EC, Vasquez YM, Li X, Kommagani R, Jiang L, Chen R, Lanz RB, Kovanci E, Gibbons WE, DeMayo FJ. 2015. Progesterone Receptor Transcriptome and Cistrome in Decidualized Human Endometrial Stromal Cells. Endocrinology 156:2239–2253. doi:10.1210/en.2014-1566

Mess A, Carter AM. 2006. Evolutionary transformations of fetal membrane characters in Eutheria with special reference to Afrotheria. J Exp Zoology Part B Mol Dev Evol 306B:140–163. doi:10.1002/jez.b.21079

Minh BQ, Schmidt HA, Chernomor O, Schrempf D, Woodhams MD, von HA. 2020. IQ-TREE 2: New Models and Efficient Methods for Phylogenetic Inference in the Genomic Era. Mol Biol Evol 37:1530–1534. doi:10.1093/molbev/msaa015

Nagy PL, Cleary ML, Brown PO, Lieb JD. 2003. Genomewide demarcation of RNA polymerase II transcription units revealed by physical fractionation of chromatin. Proc National Acad Sci 100:6364–6369. doi:10.1073/pnas.1131966100

Nguyen L-T, Schmidt HA, Haeseler A von, Minh BQ. 2015. IQ-TREE: A Fast and Effective Stochastic Algorithm for Estimating Maximum-Likelihood Phylogenies. Mol Biol Evol 32:268–274. doi:10.1093/molbev/msu300

O’Mara TA, Spurdle AB, Glubb DM, Consortium ECACECA. 2019. Analysis of Promoter-Associated Chromatin Interactions Reveals Biologically Relevant Candidate Target Genes at Endometrial Cancer Risk Loci. Cancers 11:1440. doi:10.3390/cancers11101440

Patsoukis N, Wang Q, Strauss L, Boussiotis VA. 2020. Revisiting the PD-1 pathway. Sci Adv 6:eabd2712. doi:10.1126/sciadv.abd2712

Pertea M, Kim D, Pertea GM, Leek JT, Salzberg SL. 2016. Transcript-level expression analysis of RNA-seq experiments with HISAT, StringTie and Ballgown. Nat Protoc 11:1650–1667. doi:10.1038/nprot.2016.095

Pertea M, Pertea GM, Antonescu CM, Chang T-C, Mendell JT, Salzberg SL. 2015. StringTie enables improved reconstruction of a transcriptome from RNA-seq reads. Nat Biotechnol 33:290–295. doi:10.1038/nbt.3122

Phillips JB, Abbot P, Rokas A. 2015. Is preterm birth a human-specific syndrome? Evol Medicine Public Heal 2015:136–148. doi:10.1093/emph/eov010

Pijnenborg R, Vercruysse L, Carter AM. 2011a. Deep trophoblast invasion and spiral artery remodelling in the placental bed of the lowland gorilla. Placenta 32:586–591. doi:10.1016/j.placenta.2011.05.007

Pijnenborg R, Vercruysse L, Carter AM. 2011b. Deep trophoblast invasion and spiral artery remodelling in the placental bed of the chimpanzee. Placenta 32:400–408. doi:10.1016/j.placenta.2011.02.009

Plunkett J, Doniger S, Orabona G, Morgan T, Haataja R, Hallman M, Puttonen H, Menon R, Kuczynski E, Norwitz E, Snegovskikh V, Palotie A, Peltonen L, Fellman V, DeFranco EA, Chaudhari BP, McGregor TL, McElroy JJ, Oetjens MT, Teramo K, Borecki I, Fay J, Muglia L. 2011. An Evolutionary Genomic Approach to Identify Genes Involved in Human Birth Timing. Plos Genet 7:e1001365. doi:10.1371/journal.pgen.1001365

Qiu X, Hill A, Packer J, Lin D, Ma Y-A, Trapnell C. 2017. Single-cell mRNA quantification and differential analysis with Census. Nat Methods 14:309–315. doi:10.1038/nmeth.4150

Ranzil S, Ellery S, Walker DW, Vaillancourt C, Alfaidy N, Bonnin A, Borg A, Wallace EM, Ebeling PR, Erwich JJ, Murthi P. 2019. Disrupted placental serotonin synthetic pathway and increased placental serotonin: Potential implications in the pathogenesis of human fetal growth restriction. Placenta 84:74–83. doi:10.1016/j.placenta.2019.05.012

Renfree M. 1995. Monotreme and marsupial reproduction. Reproduction Fertility Dev 7:1003– 1020. doi:10.1071/rd9951003

Renfree M, Shaw G. 2013. eLS. doi:10.1038/npg.els.0001856

Rokas A, Mesiano S, Tamam O, LaBella A, Zhang G, Muglia L. 2020. Developing a theoretical evolutionary framework to solve the mystery of parturition initiation. Elife 9:e58343. doi:10.7554/elife.58343

Rosenberg KR, Trevathan WR. 2007. An anthropological perspective on the evolutionary context of preeclampsia in humans. J Reprod Immunol 76:91–97. doi:10.1016/j.jri.2007.03.011

Rosenfeld CS. 2019. Placental serotonin signaling, pregnancy outcomes, and regulation of fetal brain development. Biol Reprod 102:532–538. doi:10.1093/biolre/ioz204

Ross LE, Grigoriadis S, Mamisashvili L, VonderPorten EH, Roerecke M, Rehm J, Dennis C-L, Koren G, Steiner M, Mousmanis P, Cheung A. 2013. Selected Pregnancy and Delivery Outcomes After Exposure to Antidepressant Medication: A Systematic Review and Meta-analysis. Jama Psychiat 70:436–443. doi:10.1001/jamapsychiatry.2013.684

Sakabe NJ, Aneas I, Knoblauch N, Sobreira DR, Clark N, Paz C, Horth C, Ziffra R, Kaur H, Liu X, Anderson R, Morrison J, Cheung VC, Grotegut C, Reddy TE, Jacobsson B, Hallman M, Teramo K, Murtha A, Kessler J, Grobman W, Zhang G, Muglia LJ, Rana S, Lynch VJ, Crawford GE, Ober C, He X, Nóbrega MA. 2020. Transcriptome and regulatory maps of decidua-derived stromal cells inform gene discovery in preterm birth. Sci Adv 6:eabc8696. doi:10.1126/sciadv.abc8696

Schmid T, Snoek LB, Fröhli E, Bent ML van der, Kammenga J, Hajnal A. 2015. Systemic Regulation of RAS/MAPK Signaling by the Serotonin Metabolite 5-HIAA. Plos Genet 11:e1005236. doi:10.1371/journal.pgen.1005236

Sharpe AH, Pauken KE. 2018. The diverse functions of the PD1 inhibitory pathway. Nat Rev Immunol 18:153–167. doi:10.1038/nri.2017.108

Sharpe AH, Wherry EJ, Ahmed R, Freeman GJ. 2007. The function of programmed cell death 1 and its ligands in regulating autoimmunity and infection. Nat Immunol 8:239–245. doi:10.1038/ni1443

Soares MJ, Varberg KM, Iqbal K. 2018. Hemochorial placentation: development, function, and adaptations. Biol Reprod 99:196–211. doi:10.1093/biolre/ioy049

Soubrier J, Steel M, Lee MSY, Sarkissian CD, Guindon S, Ho SYW, Cooper A. 2012. The Influence of Rate Heterogeneity among Sites on the Time Dependence of Molecular Rates. Mol Biol Evol 29:3345–3358. doi:10.1093/molbev/mss140

Strassmann BI. 1996. The Evolution of Endometrial Cycles and Menstruation. Q Rev Biology 71:181–220. doi:10.1086/419369

Sujan AC, Rickert ME, Öberg AS, Quinn PD, Hernández-Díaz S, Almqvist C, Lichtenstein P, Larsson H, D’Onofrio BM. 2017. Associations of Maternal Antidepressant Use During the First Trimester of Pregnancy With Preterm Birth, Small for Gestational Age, Autism Spectrum Disorder, and Attention-Deficit/Hyperactivity Disorder in Offspring. Jama 317:1553–1562. doi:10.1001/jama.2017.3413

Suryawanshi H, Morozov P, Straus A, Sahasrabudhe N, Max KEA, Garzia A, Kustagi M, Tuschl T, Williams Z. 2018. A single-cell survey of the human first-trimester placenta and decidua. Sci Adv 4:eaau4788. doi:10.1126/sciadv.aau4788

Swaggart KA, Pavlicev M, Muglia LJ. 2015. Genomics of Preterm Birth. Csh Perspect Med 5:a023127. doi:10.1101/cshperspect.a023127

Tu J, Wong C-Y. 1976. Serotonin Metabolism in Normal and Abnormal Infants during the Perinatal Period. Neonatology 29:187–193. doi:10.1159/000240863

Turco MY, Gardner L, Hughes J, Cindrova-Davies T, Gomez MJ, Farrell L, Hollinshead M, Marsh SGE, Brosens JJ, Critchley HO, Simons BD, Hemberger M, Koo B-K, Moffett A, Burton GJ. 2017. Long-term, hormone-responsive organoid cultures of human endometrium in a chemically defined medium. Nat Cell Biol 19:568–577. doi:10.1038/ncb3516

Vento-Tormo R, Efremova M, Botting RA, Turco MY, Vento-Tormo M, Meyer KB, Park J-E, Stephenson E, Polański K, Goncalves A, Gardner L, Holmqvist S, Henriksson J, Zou A, Sharkey AM, Millar B, Innes B, Wood L, Wilbrey-Clark A, Payne RP, Ivarsson MA, Lisgo S, Filby A, Rowitch DH, Bulmer JN, Wright GJ, Stubbington MJT, Haniffa M, Moffett A, Teichmann SA. 2018. Single-cell reconstruction of the early maternal–fetal interface in humans. Nature 563:347–353. doi:10.1038/s41586-018-0698-6

Wagner GP, Kin K, Lynch VJ. 2013. A model based criterion for gene expression calls using RNA-seq data. Theor Biosci 132:159–164. doi:10.1007/s12064-013-0178-3

Wagner GP, Kin K, Lynch VJ. 2012. Measurement of mRNA abundance using RNA-seq data: RPKM measure is inconsistent among samples. Theor Biosci 131:281–285. doi:10.1007/s12064-012-0162-3

Wang W, Vilella F, Alama P, Moreno I, Mignardi M, Isakova A, Pan W, Simon C, Quake SR. 2020. Single-cell transcriptomic atlas of the human endometrium during the menstrual cycle. Nat Med 26:1644–1653. doi:10.1038/s41591-020-1040-z

Wildman DE, Uddin M, Romero R, Gonzalez JM, Than NG, Murphy J, Hou Z-C, Fritz J. 2011. Spontaneous Abortion and Preterm Labor and Delivery in Nonhuman Primates: Evidence from a Captive Colony of Chimpanzees (Pan troglodytes). Plos One 6:e24509. doi:10.1371/journal.pone.0024509

Yan W, Wu F, Morser J, Wu Q. 2000. Corin, a transmembrane cardiac serine protease, acts as a pro-atrial natriuretic peptide-converting enzyme. Proc National Acad Sci 97:8525–8529. doi:10.1073/pnas.150149097

Yonkers KA, Norwitz ER, Smith MV, Lockwood CJ, Gotman N, Luchansky E, Lin H, Belanger K. 2012. Depression and Serotonin Reuptake Inhibitor Treatment as Risk Factors for Preterm Birth. Epidemiology 23:677–685. doi:10.1097/ede.0b013e31825838e9

Zhang G, Feenstra B, Bacelis J, Liu X, Muglia LM, Juodakis J, Miller DE, Litterman N, Jiang P-P, Russell L, Hinds DA, Hu Y, Weirauch MT, Chen X, Chavan AR, Wagner GP, Pavličev M, Nnamani MC, Maziarz J, Karjalainen MK, Rämet M, Sengpiel V, Geller F, Boyd HA, Palotie A, Momany A, Bedell B, Ryckman KK, Huusko JM, Forney CR, Kottyan LC, Hallman M, Teramo K, Nohr EA, Smith GD, Melbye M, Jacobsson B, Muglia LJ. 2017. Genetic Associations with Gestational Duration and Spontaneous Preterm Birth. New Engl J Medicine 377:1156–1167. doi:10.1056/nejmoa1612665

Zhang Y, Liu T, Meyer CA, Eeckhoute J, Johnson DS, Bernstein BE, Nusbaum C, Myers RM, Brown M, Li W, Liu XS. 2008. Model-based Analysis of ChIP-Seq (MACS). Genome Biol 9:R137. doi:10.1186/gb-2008-9-9-r137

